# Increased mRNA translation delays lung adenocarcinoma initiation and exposes a therapeutic vulnerability to MEK inhibitors

**DOI:** 10.1101/2025.03.19.644135

**Authors:** Luis Pardo, Madeleine Moore, Ruhi Deshmukh, Ian Powley, Joseph A. Waldron, Bjorn Kruspig, Lynn McGarry, Colin Wood, Holly Leslie, Mark Hughes, Emanuel Jeldes, June Munro, Louise Mitchell, Leah Officer-Jones, Rachel Pennie, Nigel Jamieson, David Sumpton, Douglas Strathdee, John le Quesne, Martin Bushell, Daniel L. Murphy, Jim C. Norman

## Abstract

Although protein synthesis inhibitors are being evaluated as anti-cancer agents, the dynamics of mRNA translation in early tumorigenesis are still poorly understood. We report that deletion of the mRNA-translation repressor, eIF4A2 in early KRAS-driven lung adenocarcinoma leads to a dysregulated protein synthesis landscape characterised by a strongly upregulated secretome, enlarged secretory compartments, increased oxidative metabolism and acquisition of senescence-like characteristics. Paradoxically, this overdriven protein synthesis landscape delays tumorigenesis and leads to appearance of clusters of non-proliferative, p21-positive KRAS-expressing cells in the lung. Administration of rapamycin to reduce mRNA translation suppresses senescence and restores tumorigenesis following eIF4A2 deletion. Importantly, some eIF4A2 knockout cells overcome senescence to form tumours that exhibit enhanced MAP-kinase signalling and, in contrast to eIF4A2^+/+^ lesions, these may be eradicated by administration of a MEK inhibitor. Thus, dysregulated mRNA translation exposes a potential therapeutic vulnerability in KRAS-driven lung adenocarcinoma by forcing cancer cells to rely on MEK signalling.

**Statement of significance:** The requirement for anabolism in cancers has led to the search for inhibitors of mRNA translation as anti-cancer agents. However, we report that increased rates of mRNA translation promote a senescence-like phenotype in KRAS-driven lung cancer which delays tumorigenesis and renders the resulting tumours sensitive to MEK inhibition.

## Introduction

Oncogenic events during tumour progression converge on the protein synthesis machinery to sculpt mRNA translation landscapes that drive cancer cell growth, survival and invasiveness [1, 2]. Control of mRNA translation initiation and elongation is, therefore, a key regulatory node that integrates oncogenic signalling and, consequently, has become a promising target for cancer therapeutics. Drugs targeting protein synthesis include those aimed directly at components of the mRNA translation initiation machinery (including the rocaglamides, hippuristanol or the recently approved Zotatifin, that targets the RNA helicases from the translation initiation complex eIF4As) and inhibitors of mRNA translation elongation (such as homoharringtonine) [1–3]. Furthermore, inhibitors of upstream regulators of mRNA translation, such as the mTOR inhibitors, are gaining ground as potential anti-cancer drugs [4, 5].

To date, mRNA translation inhibition strategies in cancer therapy rest on the premise that cancer cells have increased demand for protein synthesis. Therefore, inhibiting the mRNA translation machinery can compromise tumour progression with less effect on healthy cells. Indeed, the mTOR inhibitor rapamycin is well-tolerated by healthy cells but is a well-established inhibitor of tumour growth in preclinical models and humans [4, 5]. However, we have recently shown that this is not always the case. In initial stages of disease, a degree of translation inhibition may benefit the acquisition of a malignant phenotype by transformed cells, and administration of rapamycin during early-stage disease can result in increased tumorigenesis [6]. In this context, the role of the eIF4A RNA helicases is critical, as it is now becoming clear that these shape the mRNA translational programmes during tumour development. There are two main eIF4A paralogues, eIF4A1 and eIF4A2 which appear to act antagonistically [7]. eIF4A1 is an activator of mRNA translation which binds to the eIF4F complex to unwind 5’ UTR structures in the RNA thus allowing binding of the 40s ribosomal subunit and ribosome scanning to find the translational start site. eIF4A2, however, preferentially binds the Ccr4-NOT complex and participates in translation repression and mRNA decay [8, 9]. The tissue expression of eIF4A2 paralogues conforms to this view, with eIF4A1 being associated with proliferative tissues and eIF4A2 with secretory ones [6, 10]. We have recently shown that eIF4A2 restrains translation of mRNAs encoding secretory and transmembrane proteins at the endoplasmic reticulum (ER) [8], while eIF4A1 promotes proliferation in intestinal crypts by regulating translation of MYC and WNT target mRNAs [10]. Consideration of the potentially shifting mRNA translation landscapes from tumour initiation to growth and progression, and the likelihood that existing eIF4A inhibitors (such as Zotatifin) do not possess paralogue specificity, it becomes clear that the effects of these drugs should be carefully assessed using autochthonous pre-clinical models that recapitulate the tumorigenesis journey - from inception to later-stage progression - before being assessed in the clinic.

We have used an autochthonous mouse model to study the role of eIF4A1 and eIF4A2 in initiation and progression of lung adenocarcinoma (LuAd) [11]. Importantly, this model permits synchronous induction of oncogenes in combination with specific gene deletion in the lung epithelium, thus allowing us to study the effect of components of the mRNA translation machinery in various stages of LuAd initiation and progression. This approach has enabled us to demonstrate that selective restraint of mRNA translation by eIF4A2 reduces metabolic and proteostatic stress, thus allowing oncogene-expressing cells to escape oncogene-induced senescence (OIS) and proceed rapidly to establish LuAd. Incumbent in this finding is the caveat that administration of drugs to reduce mRNA translation early in tumorigenesis can ameliorate these stresses and accelerate tumour growth. Finally, we have shown that stresses imposed by deletion of *Eif4a2* can render tumour cells more dependent on MAP-kinase signalling and expose a vulnerability of tumours expressing low levels of eIF4A2 to MEK-targeting drugs.

## Results

### *Eif4A2* deletion delays tumour initiation in a transgenic mouse model of KRAS-driven lung adenocarcinoma

To investigate the role of eIF4A translational regulators in the genesis and progression of lung cancer, we chose the ‘KM’ mouse model of CRE-inducible autochthonous LuAd [11] driven by endogenously expressed activated KRAS (*Kras*^LSL-G12D/WT^) in combination with human cMYC expressed from the Rosa26 locus (Rosa26^DM.LSL-MYC/LSL-MYC^) at levels that approximate *MYC* trisomy [12]. Mutant KRAS is a well-recognised driver of lung cancer while MYC shows low-level amplification in over a third of all lung cancers [13] and strongly promotes ribosome biogenesis and protein translation [14–16]. Intranasal administration of Ad5-SPC-CRE leads to expression of KRAS^G12D^ and modestly upregulated MYC in ∼5% alveolar type II cells, the most common cell of origin in LuAd (Fig. 1A). For some experiments, we also incorporated a conditional allele of near infrared fluorescent protein (*Hprt*^LSL-iRFP^) to allow monitoring of tumour growth in living mice [17]. Following Ad5-SPC-CRE induction, KRAS^G21D^-expressing cells entered a pre-tumorigenic phase of approximately 28 days during which increased phospho-ERK was detectable in numerous cells scattered across the lung parenchyma (Fig. S1A), but these cells had not yet commenced proliferation. Transformed cells then started to proliferate to yield tumours that were detectable (by histological staining and iRFP imaging) from approximately 50 days following Ad5-SPC-CRE administration (Fig. 1A-D; S1A). These tumours continued to grow leading to a clinical endpoint with a median survival of 116 days (Fig. 1E; S1A). Some experiments were conducted in *Kras*^LSL-G12D/WT^ mice without inclusion of the MYC alleles. In this case, LuAd development was slower, and the clinical endpoint was reached with a median survival of 206 days indicating that modest upregulation of MYC is sufficient to significantly accelerate tumorigenesis, thus allowing experiments to be completed in a timely manner (Fig. S1B).

**Figure 1.**
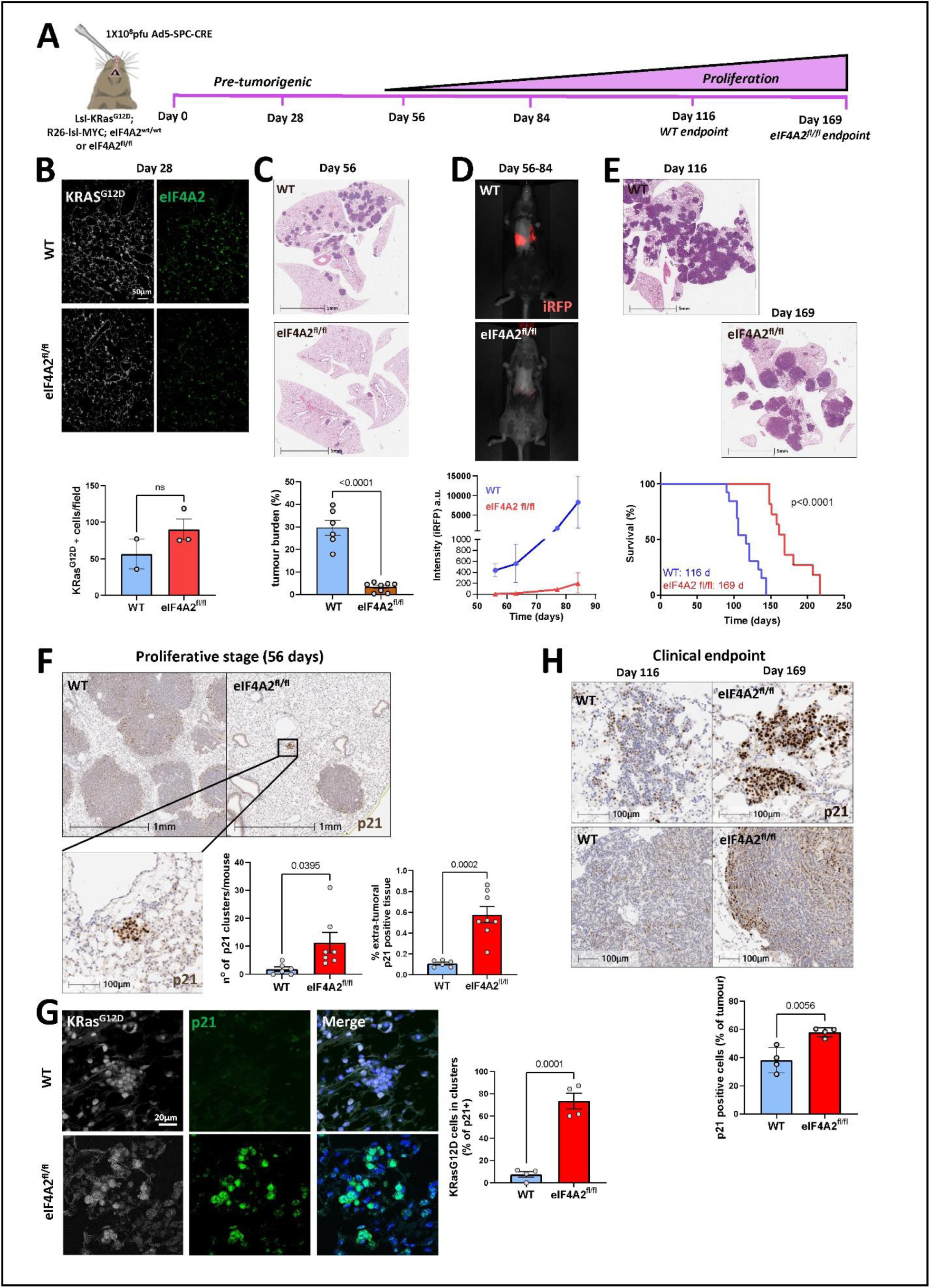
eIF4A2 deletion delays tumorigenesis of KRAS-driven lung cancer. **(A-E)** Ad5-SPC-CRE was administered intranasally to *KRAS*^LSL-G12D/WT^; *Rosa26*^LSL-MYC/LSL-MYC^ mice that were either *eIF4A2*^WT/WT^, or *eIF4A2*^fl/fl^ at a titre (1 × 10^8^ plaque forming units (pfu)/mouse) sufficient to evoke recombination in ≈5% alveolar type II cells (**A**; Day 0). Mice were sacrificed at the indicated time points **(B, C)** or at clinical endpoint **(E)** and lungs removed and fixed. Mutant KRAS^G12D^ and eIF4A2 were visualised using immunofluorescence **(B)** and lung tumour burden determined by H&E staining **(C, E)**. In **D**, Ad5-SPC-CRE was administered to *Kras*^LSL-G12D/WT^; *Rosa26*^LSL-MYC/LSL-MYC^; *Hprt*^LSL-iRFP^; mice that were either *eIF4A2*^WT/WT^ or *eIF4A2*^fl/fl^ and lung tumour progression was monitored by whole body iRFP imaging. In **(B,C)** bars are mean ± SEM, each point represents an individual mouse, and statistical test is unpaired t-test. In **(D)** graph represents mean ± SD, 2 individual mice per point. The statistical test for the Kaplan-Maier analysis in **(E)** is Logrank (Mantel-Cox) and cohort sizes were N=13 individual mice for eIF4A2^wt/wt^ (WT) and N=11 for eIF4A2^fl/fl^. **(F-H)** *Kras*^LSL-G12D/WT^; *Rosa26*^LSL-MYC/LSL-MYC^ mice that were either *eIF4A2*^WT/WT^, or *eIF4A2*^fl/fl^ were induced with Ad5-SPC-CRE as for **(A)**. Mice were sacrificed 56 days **(F, G)** following induction or at clinical endpoint **(H)** and lungs removed and fixed. p21 was visualised in the extra-tumoral regions of the lung and in lung tumours using immunohistochemistry **(F, H)**, and KRAS^G12D^ and p21 were visualised using immunofluorescence **(G)**. bars are mean ± SEM, each point represents quantification from an individual mouse, statistical test is unpaired t-test.

The eIF4A helicases, eIF4A1 and eIF4A2 are both highly expressed in lung adenocarcinoma [18]. Interestingly though, it is eIF4A2 – the eIF4A paralogue now established to act as an mRNA-translation repressor [6, 9] – that is most frequently amplified in LuAd, and this is associated with poorer overall patient survival (Fig. S2A, B). This suggested the possibility that an inhibitor of mRNA translation might be a driver of cancer aggressiveness and prompted us to study the role of eIF4A2 in the establishment and progression of LuAd. eIF4A2 was upregulated in KRAS^G21D^-expressing cells during the pre-tumorigenic phase (i.e. up to 28 days following Ad5-SPC-CRE administration) of LuAd (Fig. 1B; S1C). However, as tumours exited the pre-tumorigenic phase and started to grow, eIF4A2 levels declined (particularly in the more proliferative tumour periphery), consistent with a role for this endogenous inhibitor of mRNA translation in initiation events occurring in early LuAd tumorigenesis and before transformed cells start to proliferate (Fig. S1D, left panels). To test this, we crossed KM mice with mice harbouring homozygous floxed alleles of *Eif4a2*. This led to efficient deletion of *Eif4a2* in KRAS-expressing cells when stained 28 days following Ad5-SPC-CRE administration (Fig. 1B). Importantly, the ability of these KRAS-expressing tumour-initiating cells to commence proliferation and form tumours was severely retarded by *Eif4a2* deletion. Indeed, in KM*-Eif4a2*^fl/fl^ mice tumour formation was largely undetectable (using either histological or iRFP imaging approaches) up to 85 days following Ad5-SPC-CRE administration (Fig. 1C, D). This retardation of tumour growth led to a significant extension of median survival (from 116 to 169 days); with these animals exhibiting fewer (but larger) tumours at clinical endpoint than KM*-Eif4a2*^wt/wt^ (WT) mice (Fig. 1E). Importantly, the tumours that did grow in KM-*Eif4a2*^fl/fl^ mice did not express eIF4A2 and, therefore, were not ‘escapers’ that had evaded efficient recombination of the floxed *Eif4a2* alleles (Fig. S1D). By contrast, tumours arising in KM*-Eif4a1*^fl/fl^ mice still exhibited eIF4A1 expression and had, therefore, originated from transformed cells that escaped recombination of the floxed *Eif4a1* allele (Fig. S1D). Finally, the ability of conditional eIF4A2 deletion to increase survival was even more marked in the absence of MYC overexpression. Indeed, *Eif4a2* deletion extended median survival from 206 days to a value that exceeded 337 days; with 9 out of the 14 *Kras*^LSL-G12D/WT^; *Eif4a2*^fl/fl^ mice displaying no signs of disease 375 days following Ad5-SPC-CRE administration (Fig. S1B). However, because the constraints of our animal licence precluded continuation of this experiment beyond 375 days following Ad5-SPC-CRE administration, we were unable to obtain a quantitative measure (median survival) of the ability of *Eif4a2* deletion to extend survival from KRAS-driven LuAd in the absence of MYC overexpression.

Oncogene-induced senescence (OIS) is thought to suppress tumour formation by arresting proliferation in cells that have acquired potentially oncogenic mutations [19, 20]. Thus, the capacity of mutated cells to override OIS can enable tumour initiation. We hypothesised that eIF4A2, possibly by restraining mRNA translation, might influence the ability of KRAS^G12D^ and its downstream signalling to promote OIS and/or the ability of oncogene-expressing cells to override senescence programmes and commence proliferation. As outlined above, *Eif4a2* deletion delays transition from a non-proliferative pre-tumorigenic state to one in which there is rapid tumour growth. Whilst extensive tumour formation (≈30% of lung area encompassed by tumour tissue) was seen in KM-*Eif4a2*^wt/wt^ (WT) mice 56 days following KRAS/MYC activation, few tumours were present in the lungs of KM-*Eif4a2*^fl/fl^ mice at this time point (Fig. 1C). Instead, we observed clusters of KRAS^G12D^-expressing lung epithelial cells the majority of which were positive for the senescence marker, p21. Extra-tumoral clusters of KRAS^G12D^-expressing cells were much less frequent in KM-*Eif4a2*^wt/wt^ (WT) mice and contained fewer p21-positive cells (Fig. 1F,G). Interestingly, extra-tumoral clusters of p21-positive cells were still visible in the lung of KM-*Eif4a2*^fl/fl^ mice at clinical endpoint, and the tumours that did grow in these animals had an increased proportion of p21-expressing cells than the corresponding ‘endpoint’ tumours in KM-*Eif4a2*^wt/wt^ mice (Fig 1H).

### *Eif4a2* deletion evokes a senescence-like phenotype in lung tumour cells

Cellular senescence – whether evoked by oncogenes (OIS), toxic insults or cell ageing - is characterised by acquisition of phenotypic behaviours, termed ‘senescence hallmarks’, which can include: i) exit from the cell cycle; ii) altered nuclear morphology; iii) expression of proteins associated with cell cycle arrest (e.g. p21 and p16); iv) lysosomal expansion (as indicated by β-galactosidase expression) and; v) acquisition of a highly secretory state characterised by expansion of biosynthetic secretory compartments and copious production of a secretome, termed the senescence-associated secretory phenotype (SASP) [21]. Acquisition of OIS is also reported to be associated with marked alterations to cellular metabolism characterised by activation of pyruvate dehydrogenase driving flux of glycolytic carbons into the Krebs cycle, and increased oxidative phosphorylation (oxPHOS) and generation of reactive oxygen species (ROS) [22]. To investigate the role of eIF4A2 and the mRNA translation machinery in these cellular processes, we grew tumour cells from the lung tumours of a KM mouse. We then used CRISPR to delete *Eif4a2* from KM cells (termed KM^4A2cr^) and generated non-targeting CRISPR control KM cells (termed KM^NTC^) (Fig. 2A, B). KM^NTC^ and KM^4A2cr^ cells grew at similar rates in culture, and both displayed clonogenic potential when adherent to plastic dishes (Fig. 2C). However, KM^4A2cr^ cells had significantly reduced ability to form spheroids when grown in low attachment conditions without serum (Fig. 2D) indicating altered tumorigenic properties. Consistent with this, KM^4A2cr^ cells displayed several phenotypic characteristics associated with cellular senescence including: i) upregulated expression of nuclear p21 (Fig2E); ii) increased nuclear area (Fig. 2G); iii) increased β-galactosidase activity (Fig. 2H); iv) expansion of the Golgi apparatus (Fig. 2F); and v) increased O_2_ consumption (indicating upregulated oxPHOS) (Fig. 2I) and elevated ROS generation (Fig. 2J).

**Figure 2.**
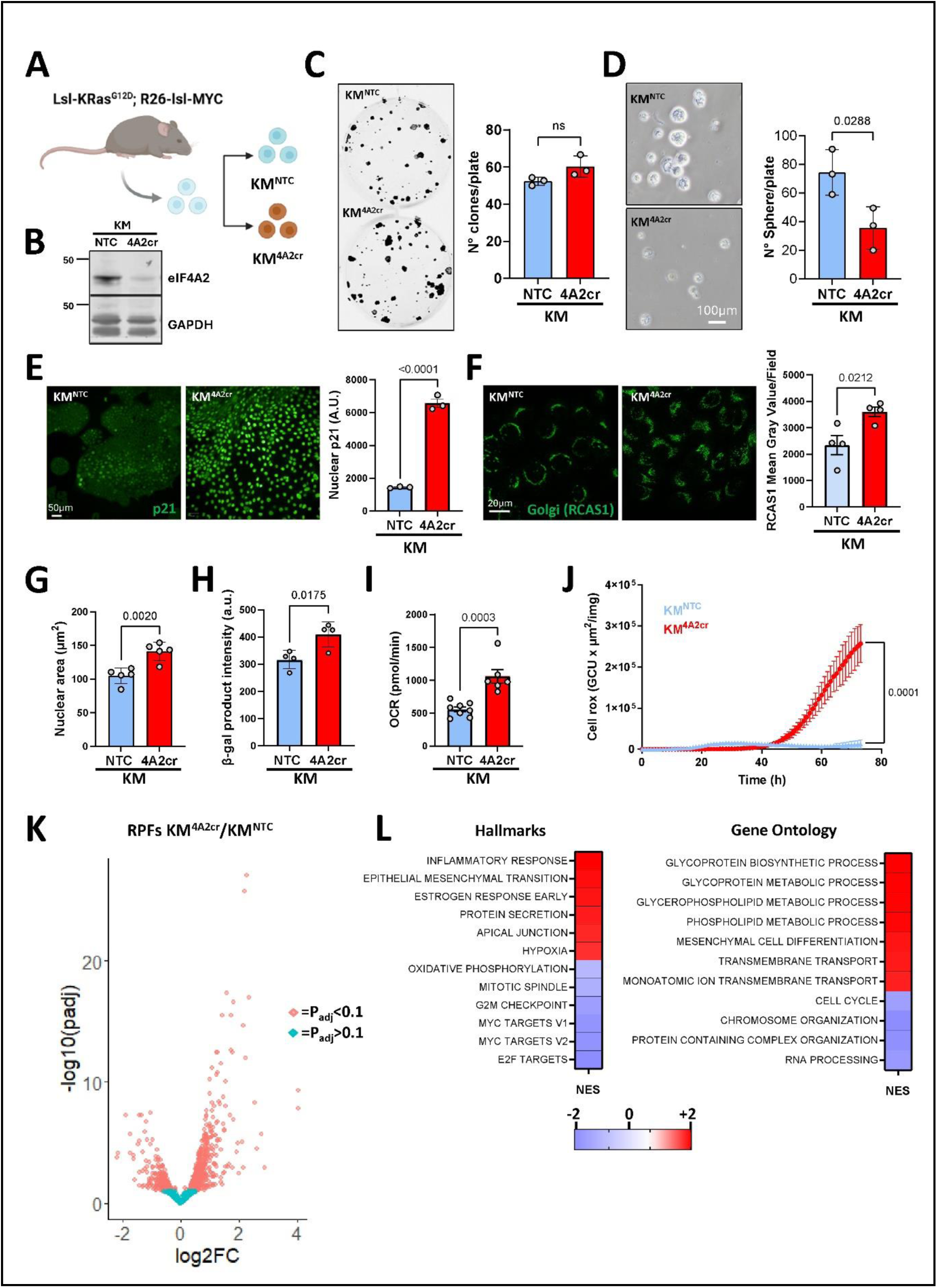
eIF4A2 deletion evokes a senescence-like phenotype in cells from KRAS-driven lung cancer. **(A-B)** *KRAS*^LSL-G12D/WT^; *Rosa26*^LSL-MYC/LSL-MYC^ (KM) mice were induced intranasally with Ad5-SPC-CRE. Lung tumours were resected from these mice and cell lines established from these (KM cells). KM cells were transduced with lentiviral vectors bearing guide RNAs recognising non-targeting sequences (NTC) or sequences in eIF4A2 (4A2cr), and eIF4A2 expression in the resulting KM^NTC^ and KM^4A2cr^ cells was determined by Western blot using GAPDH as loading control (B). **(C, D)** KM^NTC^ or KM^4A2cr^ cells were plated at low density onto plastic dishes in serum containing medium (C) or into low adhesion wells in serum-free medium (D). The resulting number of colonies (C) and tumour spheres (D) respectively were then determined. Bars are mean ± SEM, N=3 individual experiments, statistical test is unpaired t-test. **(E - J)** KM^NTC^ or KM^4A2cr^ cells were plated onto plastic surfaces and allowed to grow for 48 hr. p21 expression (E), area of the Golgi (F; RCAS1 is a cis-Golgi marker), nuclear area (G) and β-galactosidase activity (H), were visualised and quantified using immunofluorescence. Graph bars are mean ± SEM of N=3-5 individual experiments, statistical test is unpaired t-test. Oxygen consumption rate (I; OCR) and cellular reactive oxygen species (J: cell rox) were determined using Seahorse XF analysis and the CellROX reagent respectively. Graph in I is representative of n=3 individual experiments. Bars represent mean ± SEM of 5 technical replicates analyzed by unpaired t-test. Graph in J is mean ± SEM of N=6 independent experiments analyzed by linear regression, **(K, L)** KM^NTC^ or KM^4A2cr^ cells were grown for 48 hr on plastic dishes and lysates subjected to ribosome footprinting analysis in which the regions of mRNA which were protected from nuclease digestion by ribosome occupancy were sequenced. Volcano plot shows and increased number of ribosome-protected fragments (RPFs) in KM^4A2cr^ (positive log2FC values) than in KM^NTC^ (negative log2FC values). The heatmap displays categories of RPFs significantly enriched (red indicates positive normalised enrichment score (NES)) or depleted (blue indicates negative NES) in KM^4A2cr^ cells categorised using the ‘hallmarks’ (left column) and ‘gene ontology biological process’ (right column) tools. Data was obtained from N=4 individual experiments.

Ribosome footprinting determines the positions of ribosomes on mRNAs at the whole-genome level and, therefore, can provide a quantitative indication of which mRNAs are being translated [23]. We, therefore, used ribosome footprinting to characterise how deletion of *Eif4a2* influences the mRNA translation landscapes of KM cells. Given eIF4A2’s emerging role as a repressor of mRNA translation initiation, it was unsurprising that KM^4A2cr^ cells exhibited a net increase in ribosome occupancy (Fig. 2K; S3). Enrichment analysis of the ribosome-occupied mRNAs indicated that *Eif4a2* deletion increased translation of secretory and transmembrane proteins, particularly those associated with post-translational processing in the ER and Golgi. Indeed, *Eif4a2* deletion drove significant increases in the normalised enrichment score for categories of mRNAs (encoding proteins involved in synthesis and processing of secretory glycoproteins and secreted components of the inflammatory response (Fig. 2L). Conversely, mRNAs encoding proteins driving cell cycle progression are amongst the (smaller) cohort of genes whose ribosome occupancy was decreased following *Eif4a2* deletion. These data are consistent with the view that eIF4A2 contributes to the suppression of senescence-like features by restraining synthesis of proteins responsible for secretome production and promoting translation of mRNAs encoding cell cycle regulators.

### eIF4A2 influences cellular metabolism via mRNA translation-dependent and - independent routes

Protein synthesis is thought to put significant demands on the cell’s energy metabolism and ATP-generating capacity [24, 25], suggesting that the increased mRNA translation of secretory proteins caused by *Eif4a2* deletion may contribute to increased energetic expenditure in KM^4A2cr^ cells. We, therefore, used liquid chromatography-mass spectrometry (LC-MS) to determine the consequences of deleting *Eif4a2* on the metabolic landscape of KM cells. Firstly, we measured exchange of key metabolic fuels/metabolites between the cells and the surrounding medium. *Eif4a2* deletion significantly elevated uptake of glucose, glutamine and pyruvate, and increased lactate release and O_2_ consumption (Fig. 3A, B), indicating that *Eif4a2* deletion elevates flux through glycolysis and increases oxPHOS. To determine which of these events were driven by the energetic demands of increased protein synthesis occurring following *Eif4a2* deletion, we treated KM^NTC^ and KM^4A2cr^ cells with drugs established to reduce mRNA translation initiation. Rapamycin, a drug with well-established pharmacokinetics and capacity to target the TORC1-4EBP1 and TORC1-S6K axes (and thus oppose mRNA translation) in vivo and in vitro, significantly opposed the ability of *Eif4a2* deletion to increase pyruvate uptake and O_2_ consumption but altered neither glucose and glutamine uptake nor lactate release (Fig. 3A, B). Moreover, eFT226, a potent and selective inhibitor of eIF4A paralogues [26], significantly opposed the ability of *Eif4a2* deletion to increase O_2_ consumption (Fig. 3B). These data suggested that eIF4A2 may control several metabolic processes, not all of which are consequent upon its influence on mRNA translation/protein synthesis. We then conducted a more in-depth screen to characterise altered intracellular metabolite levels following *Eif4a2* deletion, and to identify which of these changes are reversed by rapamycin addition. This indicated that *Eif4a2* deletion suppressed level of sugars (including glucose-6-phosphate, fructose-1,6-bisphosphate and glyceraldehyde-3-phosphate) in the proximal part of glycolysis, but significantly increased components of the pentose-phosphate pathway (PPP), the feed from glycolysis into the Krebs cycle (pyruvate and acetyl-CoA) and Krebs cycle metabolites themselves (Fig. S4). Importantly, compounds downstream of the PPP, many of which are players in nucleotide synthesis and salvage, are some of the most significantly upregulated metabolites following *Eif4a2* deletion (Fig. S5). Indeed, we highlight that *Eif4a2* deleted cells exhibit >10-fold increase in hypoxanthine with no corresponding alteration in the levels of xanthine and urate, indicating a hugely increased potential for nucleotide salvage and/or a shutdown of nucleotide catabolism (Fig. S5A). Importantly, very few of the alterations to intracellular metabolite levels driven by *Eif4a2* deletion were significantly reversed by addition of rapamycin. Taken together, these data suggests that eIF4A2 influences metabolism in ways that are both dependent and independent of its control of mRNA translation; with eIF4A2’s ability to limit pyruvate uptake and oxPHOS being linked to its function as a restrainer of protein synthesis, whilst *Eif4a2*-deletion-driven upregulation of nucleotide synthesis occurred regardless of mRNA translation initiation rates.

**Figure 3.**
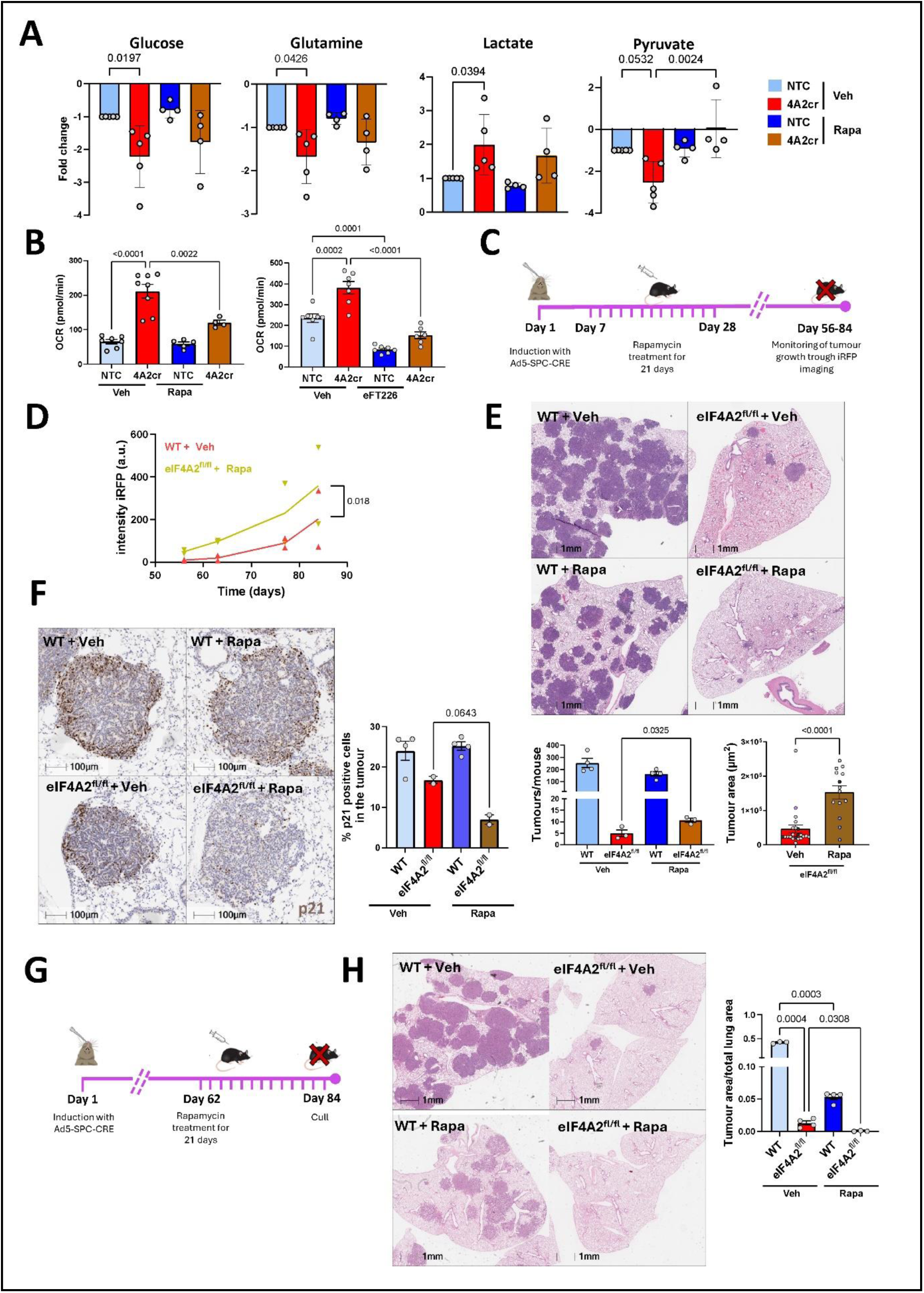
Rapamycin administration rescues increased pyruvate uptake and oxygen consumption and restores tumour growth in eIF4A2-deleted lung cancer. **(A)** KM^NTC^ or KM^4A2cr^ cells were plated onto six-well dishes and allowed to adhere overnight. Adherent cells were incubated at 37°C for 24 hr in the presence and absence of rapamycin (1 μM) or vehicle control (Veh.), medium was then aspirated, and levels of the indicated metabolites in the conditioned medium were determined using LC-MS-based metabolomics. Data are expressed as the fold-change difference between metabolite peak areas detected in cell-conditioned medium and those in medium incubated at 37°C in the absence of cells (normalised to KM^NTC^ cells). Thus, positive and negative values indicate production/release and consumption respectively during the 24 hr period. Bars are mean ± SEM, N=5 (Veh.) and N=4 (Rapa.) individual experiments. **(B)** KM^NTC^ or KM^4A2cr^ cells were allowed to adhere overnight and then treated with rapamycin (1 μM), efT226 (1 μM) or vehicle control (Veh.) as indicated for 1 hr. The oxygen consumption rate (OCR) was then determined using Seahorse XF analysis. Bars are mean ± SEM, n=7 (Veh.), n=4 (Rapa.) and n=7 (efT226) technical replicates. Graphs are representative of N=3 individual experiments. **(F-H)** *KRAS*^LSL-G12D/WT^; *Rosa26*^LSL-MYC/LSL-MYC^ (KM) mice that were either *eIF4A2*^WT/WT^, or *eIF4A2*^fl/fl^ were induced with Ad5-SPC-CRE as for figure 1A. Rapamycin (10mg/kg) or vehicle control was administered daily (by *i.p.* injection) for a 3-week period from either 7 **(C)** or 62 **(G)** days following Ad5-SPC-CRE administration. Mice were sacrificed 84 days following Ad5-SPC-CRE administration, and lungs were removed and fixed **(E, F, H)**. In **(D)**, KM mice carrying *Hprt*^LSL-iRFP^ allele were used, rapamycin administered as for **(C)** and monitoring was by whole body iRFP imaging. Graph represents mean ± SEM of 2 individual mice per condition analysed by linear regression. Lung tumour burden was determined by H&E staining **(E, H)** and p21 visualised and quantified using immunohistochemistry **(F)**. Bars are mean ± SEM, each dot represents an individual mouse, statistical test is one-way ANOVA. In the right graph of **(E)**, bars are mean ± SEM of individual tumours (represented by dots) from N=3 mice per group (each mouse tumours are colour coded in the graph), statistical test is unpaired t-test.

### Rapamycin administration allows senescence exit and restores tumour growth to *Eif4a2* knockout LuAd

We have previously reported that *Eif4a2* deletion in oncogene-expressing hepatocytes delays liver tumorigenesis. This is because the aberrantly elevated mRNA translation in these cells upregulates trafficking of integrins to lysosomes which, in turn, reduces extracellular matrix (ECM) deposition. Importantly, this delay to liver tumorigenesis is obviated by administration of rapamycin to reduce mRNA translation and restore ECM deposition following oncogene-activation [6]. We, therefore, reasoned that the increased oxidative metabolism (pyruvate and O_2_ consumption) evoked by *Eif4a2* deletion in lung epithelial cells might constitute a metabolic stress that helps to maintain OIS. To test this, we treated *Hprt*^LSL-iRFP^ positive KM mice that were either *Eif4a2*^wt/wt^ (WT) or *Eif4a2*^fl/fl^ with rapamycin (by daily intra-peritoneal injection) 7 days following Ad5-SPC-CRE administration and maintained this treatment for 21 days (Fig. 3C). This protocol was designed to reduce the increased mRNA translation, that we hypothesise to be maintaining senescence, occurring up to 28 days following oncogene activation, but then to subsequently allow protein synthesis for tumour growth. We monitored tumour growth by iRFP imaging between 56 and 84 days following Ad5-SPC-CRE administration, then sacrificed the animals to assess tumour burden and p21 status. Our results indicate that rapamycin treatment restored growth to *Eif4a2* knockout tumours (Fig. 3D) leading to a significantly increased number and burden of tumours in KM*-Eif4a2*^fl/fl^ mice (Fig. 3E) and reduced expression of p21 (indicating senescence exit) (Fig. 3F) when assessed 84 days following Ad5-SPC-CRE administration. We also conducted experiments in which we treated mice with rapamycin between 62 – 84 days following Ad5-SPC-CRE administration; a time window after tumours have exited their non-proliferative/senescent-like phase (Fig. 3G). This indicated that rapamycin can restrict tumour growth, as would be expected for an agent which reduces protein synthesis, when administered once tumours have commenced proliferation (Fig. 3H).

Taken together, these data indicate that inhibition of mRNA translation, when enacted following oncogene activation but prior to establishment of a proliferative phenotype, facilitates escape from OIS and promotes tumorigenesis.

### *Eif4a2* knockout tumours upregulate MAP-kinase signalling to evade OIS

Analysis of lungs of KM*-Eif4a2*^fl/fl^ mice at clinical endpoint indicates that, despite a minority (≈10%) of ‘escaper’ tumours (in which recombination of the floxed alleles was incomplete and eIF4A2 expression was detectable), genuinely eIF4A2-negative tumours (Fig. S6A) were able to progress, but with a delay of ≈30 days. Following emergence from the initial growth delay, eIF4A2 deficient tumours grew at a rate comparable to their eIF4A2-expressing counterparts (Fig. S6B) and, unlike tumours from WT mice, were strongly Ki67-positive at clinical endpoint (Fig. S6C). Coupling multiplex imaging with spatial transcriptomics enabled comparison of gene expression profiles of pan cytokeratin (pan CK)-positive endpoint tumours which were either eIF4A2-positive or negative and thus provided insight into how tumours circumnavigate the senescence-like state evoked by Eif4a2 deletion and acquire the ability to proliferate (Fig. 4A). Profiling of *Eif4a2* deficient tumours indicated a more aggressive behaviour compared with their *Eif4a2*^wt/wt^ (WT) counterparts, showing upregulation of proliferative and pro-survival pathways like JAK-STAT, mTORC1 and KRAS, together with increases in TGFβ signalling and markers of EMT (Fig. 4B). Furthermore, *Eif4a2* deleted tumours displayed signatures consistent with increased metabolism (glycolysis) and dysregulated synthesis of secretory proteins (unfolded protein response) consistent with what we have described previously in KM^4A2cr^ cells. To further substantiate these observations, we resected endpoint tumours from KM-*Eif4a2*^wt/wt^ (WT) and KM-*Eif4a2*^fl/fl^ mice and grew them in culture (Fig. 4C). The resulting cell lines (termed KM^A2+/+^ and KM^A2−/−^ cells) recapitulated the aggressiveness of endpoint tumours. Indeed, KM^A2−/−^ cells showed increased growth in colony formation and spheroid formation assays (Fig. S7A, B) despite maintaining some senescent features including p21 expression and ROS production (Fig. S7C, D). We used these cells to assess signalling downstream of KRAS by Western blotting and observed that KM^A2−/−^ cells exhibited marked upregulation of phospho-ERK1/2, indicating that the upregulated ‘KRAS signalling’ observed in *Eif4a2* knockout tumours was manifested as a substantial increase in MAP-kinase activity (Fig. 4D). We then confirmed that phospho-ERK levels were also elevated in KM-*Eif4a2*^fl/fl^ mice during all stages of tumour development (Fig. S6D). Conversely, phosphorylation of AKT, an indicator of increased PI-3 kinase signalling (which can also occur downstream of KRAS) was strongly suppressed in KM^A2−/−^ cells (Fig. 4D). Furthermore, KM^A2−/−^ cells exhibited increased phosphorylation of the ER-stress sensor, PERK and its downstream effector ATF4, indicating activation of the unfolded protein response as also identified in the spatial transcriptomic analysis of endpoint tumours (Fig. S7E). Taken together, these observations indicate KRAS-driven tumours adapt to loss of eIF4A2 by upregulating MAP-kinase and downregulating PI-3-kinase signalling. To determine whether reduced levels of eIF4A2 might predicate a similar signalling switch in lung cancer patients we used publicly available data from TCGA [27, 28] and the Lattice-A cohort of human lung cancer [29]. RNAseq data extracted from c-Bioportal showed that *EIF4A2* expression correlates positively with *PI3K* and negatively with *ERK1* in a cohort of 1053 lung cancer patients (Fig. S6E). We then interrogated the Lattice-A tissue microarray using a multiplex imaging approach and observed that relative activities of MAP-kinase and PI-3 kinase read-outs (phospho-ERK/phospho-AKT ratio) inversely correlated with increasing levels of eIF4A2 (but not eIF4A1), indicating that human NSCLC with low eIF4A2 expression exhibits increased dependency on MAP-kinase signalling (Fig 4. E, F).

**Figure 4.**
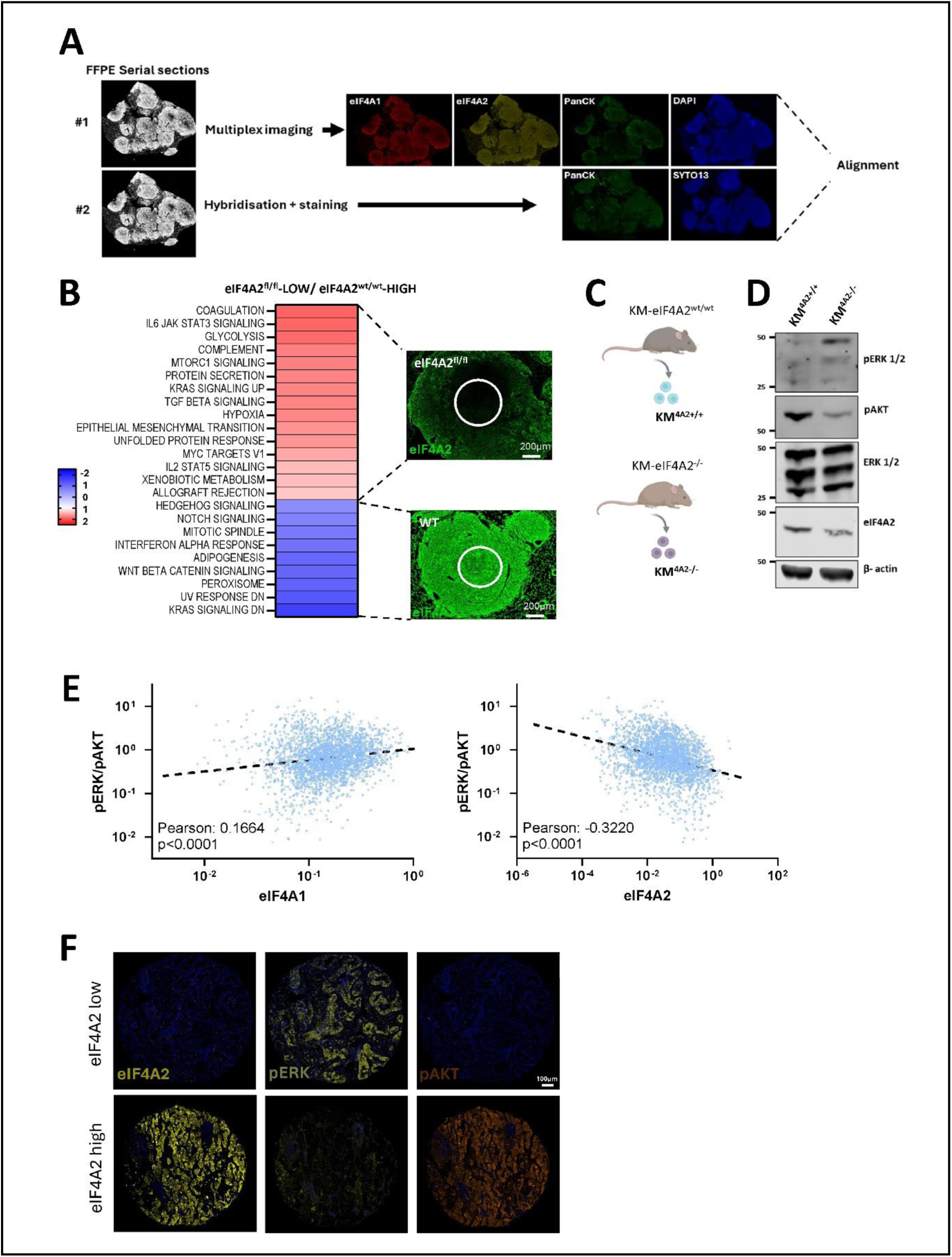
eIF4A2 knockout tumours upregulate MAP-kinase signalling. **(A-B)** *KRAS*^LSL-G12D/WT^; *Rosa26*^LSL-MYC/LSL-MYC^ (KM) mice that were either *eIF4A2*^WT/WT^, or *eIF4A2*^fl/fl^ were induced with Ad5-SPC-CRE as for figure 1A. Lungs were removed at clinical endpoint, formalin-fixed/paraffin-embedded (FFPE) and sectioned. Consecutive serial sections were subjected to multiplex imaging using antibodies recognising eIF4A1, eIF4A2, cytokeratins (pan-CK) and nuclear staining (DAPI) (#1) or spatial transcriptomics hybridisation with pan-CK and nuclear staining (SYTO13) to enable alignment (#2). The transcriptomic signatures in regions of interest (ROIs) representing the eIF4A2-negative tumours of eIF4A2^fl/fl^ mice and the eIF4A2-positive tumours of eIF4A2^WT/WT^ mice were then compared **(B)**. The heatmap displays the ‘hallmark’ gene expression signatures that were upregulated (red) and downregulated (blue) in eIF4A2-negative tumours **(B)**. **(C-D)** Tumours were resected at clinical endpoint from KM mice that were either *eIF4A2*^WT/WT^, or *eIF4A2*^fl/fl^ and cell lines established from these were termed KM^A2+/+^ and KM^A2−/−^ respectively **(C)**. Activity of MAP- and PI-3 kinase signalling in KM^A2+/+^ and KM^A2−/−^ cells was then assessed using Western blotting with antibodies recognising phospho-ERK and phospho-AKT **(D)**. β-actin was used as a loading control for eIF4A2 and total ERK as a loading control for pERK and pAKT. **(E-F)** Levels of eIF4A1, eIF4A2, phospho-ERK and phospho-AKT were quantified using multiplex immunofluorescence in tumour cores from the Lattice-A cohort of human lung cancer. The graphs indicate the relationships between the ratio of MAP- to PI-3 kinase signalling (phospho-ERK/phospho-AKT) expressed as a function of levels of eIF4A1 (left graph) or eIF4A2 (right graph) **(E)**. Each blue dot represents an individual tumour core (N=994 patients, 3 cores/patient) and the dotted line denotes analysis by linear regression, regression coefficients and p-values are displayed. Representative examples of tumour cores exhibiting high and low eIF4A2 levels and corresponding phospho-ERK and phospho-AKT expression are displayed **(F)**.

### *Eif4a2* deletion exposes a therapeutic vulnerability to MEK inhibition

Inhibition of MAP-kinases is currently proving effective in targeting solid tumours, and the MEK1/2 inhibitor, Trametinib is now an approved therapy for melanoma [30]. However, intrinsic resistance to Trametinib has limited its effectiveness in treating lung cancer [31]. We reasoned that the increased MAP-kinase signalling we observe in *Eif4a2* deleted LuAd might expose a potential therapeutic vulnerability to Trametinib in tumours with low eIF4A2 levels. Initially, we measured proliferation of KM^A2+/+^ and KM^A2−/−^ cells in the presence and absence of Trametinib. This indicated that proliferation of KM^A2−/−^ cells was significantly reduced by Trametinib, whilst KM^A2+/+^ cells were unaffected (Fig. 5A, B). To determine whether Trametinib might oppose growth of *Eif4a2*-deleted LuAd in vivo, we treated KM mice that were either *Eif4a2*^wt/wt^ (WT) or *Eif4a2*^fl/fl^ with Trametinib from 28 to 84 days following Ad5-SPC-CRE administration (Fig. 5C). Following this, mice were sacrificed, and the number of tumours was determined. This showed that Trametinib strongly suppressed growth of *Eif4a2*-deleted tumours leading to a profound reduction in the number of tumours in KM*-Eif4a2*^fl/fl^ mice, whilst tumour growth was unaffected by Trametinib in KM*-Eif4a2*^wt/wt^ (WT) mice (Fig. 5D, E). Taken together, these data indicate that reduced eIF4A2 imposes pressures leading to selection of cells with reduced PI-3 kinase and upregulated MAP-kinase signalling which can evade OIS and grow aggressively, but that are exquisitely sensitive to Trametinib.

**Figure 5.**
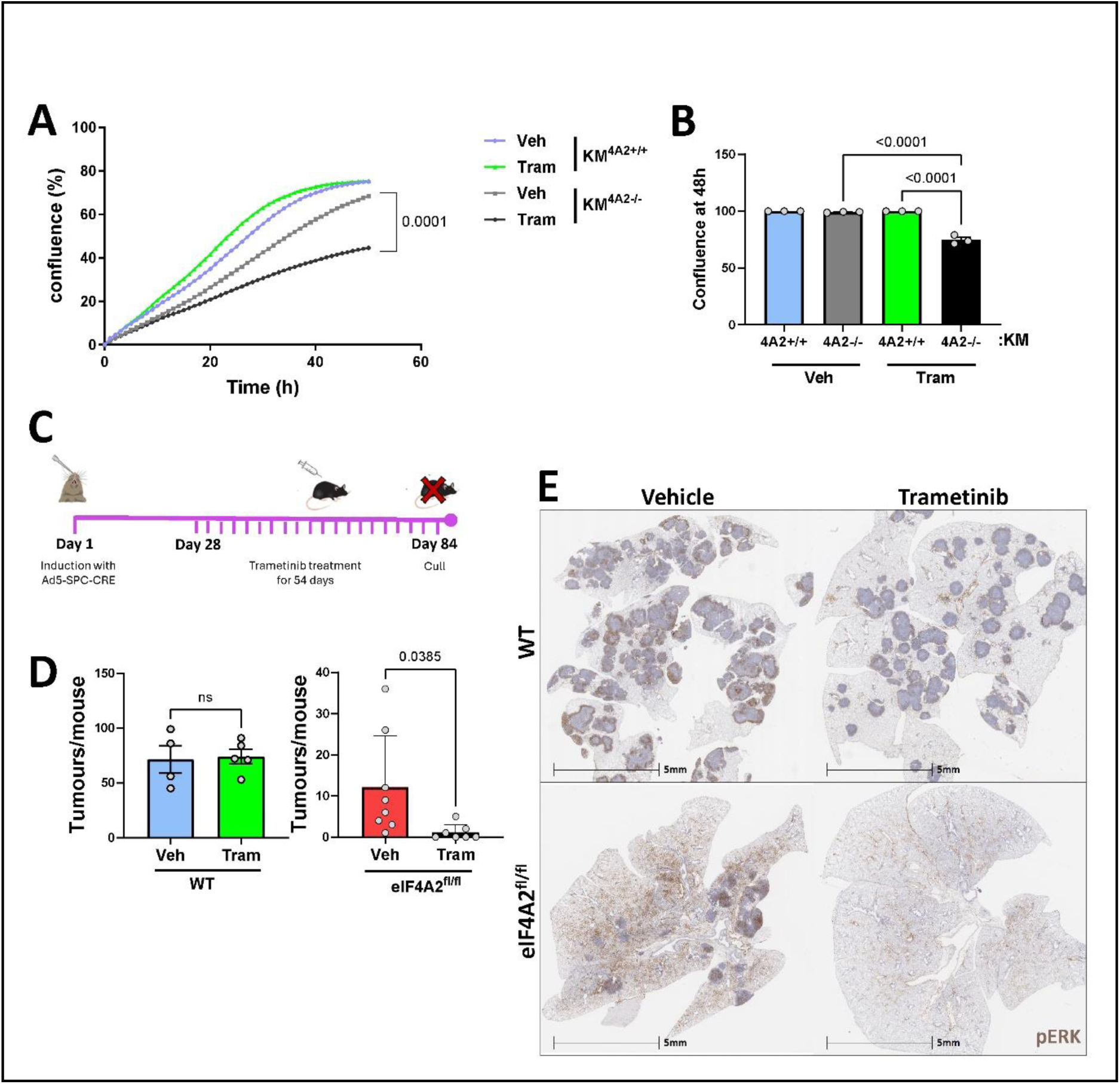
eIF4A2 deletion exposes a therapeutic vulnerability to MEK inhibition. **(A-B)** KM^2A+/+^ or KM^2A−/−^ cells were plated in the presence or absence of Trametinib (Tram.; 100 nM) or vehicle control (Veh.) and cell growth monitored over a 48 hr period **(A)**. Cell confluence was determined 48 hr following plating **(B)**, Graphs represent mean ± SEM, N=3 individual experiments. Statistical tests are linear regression in **(A)** and one way ANOVA in **(B)**. **(C-E)** *KRAS*^LSL-G12D/WT^; *Rosa26*^LSL-MYC/LSL-MYC^ (KM) mice that were either *eIF4A2*^WT/WT^, or *eIF4A2*^fl/fl^ were induced with Ad5-SPC-CRE as for figure 1A. Trametinib (1mg/kg) or vehicle control was administered daily (by *i.p.* injection) from 28 days following Ad5-SPC-CRE administration. Mice were sacrificed 84 days following Ad5-SPC-CRE administration, and lungs were removed and fixed. Lung tumour number was determined by H&E staining **(D)** and phospho-ERK visualised by immunohistochemistry **(E)**. Values are mean ± SEM, each dot represents an individual mouse, statistical test is unpaired t-test.

## Discussion

In this study we highlight an important role for a restrainer of mRNA translation as cancer cells navigate the earlier stages of tumour initiation and establishment in the lung. Contrary to an established paradigm that asserts the need for continuous activation of protein synthesis to support tumour growth, our study demonstrates that during the early stages of tumorigenesis, translation of mRNAs encoding secretory proteins is restrained by the mRNA translational repressor, eIF4A2. Such restraint minimises metabolic and proteostatic stress, and suppresses senescence-like behaviours, allowing oncogene transformed cells to proliferate and tumours to progress without delay. Despite this delay, eIF4A2 knockout cells can, with time, circumnavigate the block to tumorigenesis imposed by unrestrained mRNA translation, and they achieve this by upregulating MAP-kinase signalling. In doing this, lung cancer cells acquire a vulnerability to MEK inhibitors that potentially may be exploited to target lung tumours with low expression of eIF4A2.

Despite sharing more than 80% homology at the amino acid level, the RNA helicases eIF4A1 and eIF4A2 play distinct roles in protein synthesis. eIF4A1, which activates mRNA translation initiation, predominates in proliferative tissues such as intestinal crypts [10]. Indeed, eIF4A1 commands a translational programme comprising oncogenic drivers, such as components of the Wnt signalling cascade, which drives proliferation of intestinal cells following deletion of the APC tumour suppressor [10]. Conversely, eIF4A2 (via association with the Ccr4-NOT complex) reduces mRNA translation and is more widely expressed in less proliferative, secretory cells including those of the intestinal villi [6, 10]. eIF4A paralogue expression patterns shift during progression of NSCLC in a manner that is consistent with eIF4A1 and eIF4A2 being activator and restrainer of mRNA translation respectively. eIF4A2 expression is high in non-proliferative, senescent-like cells which characterise early KRAS-driven lung tumorigenesis, whereas eIF4A2 levels decline with tumour progression and eIF4A1 expression predominates as lung tumour cells proliferate. Consistently, reducing mRNA translation by administration of rapamycin once eIF4A2 levels have declined inhibits tumour growth, indicating that it is primarily eIF4A1’s proliferative translational programme that is being targeted when protein synthesis inhibitors are successful in cancer therapy. However, our observations that rapamycin can promote tumour initiation when administered early in tumorigenesis – when a secretory phenotype prevails over a proliferative one - highlights that targeting secretory translatomes associated with eIF4A2 may be counterproductive in cancer therapy.

Adoption of OIS is often a primary response to acquisition of mutated oncogenes, and evidence suggests that this may be an important tumour suppression mechanism in several tissues. Mutation of KRAS in the pancreatic ductal epithelium leads to accumulation of pancreatic intraepithelial neoplasia (PanINs) which are lesions with senescence-like characteristics and that can produce a SASP-like secretome [32, 33]. Similarly, mutation of NRAS or BRAF in the skin promotes formation of nevi containing senescent melanocytes with SASP-generating capacity [34]. The ability of the mutated cells to bypass this senescence-like state and to start proliferating permits progression of PDAC and melanoma respectively. Whilst cells in deep senescence undergo extensive chromatin cleavage resulting in stable cell cycle arrest, more circumspect adoption of senescence-like behaviours, which can occur without commitment to deep-senescence, may also oppose cell cycle progression, but in a reversible manner [35]. Therefore, it is thought that if activated oncogenes can evoke sufficient senescent hallmarks to slow growth and produce SASP to attract immune cells, thus providing opportunities for clearance of mutated cells, adoption of senescence hallmarks may constitute an important tumour-suppression mechanism [19, 20, 36]. Nevertheless, if the senescence programme is insufficiently deep, and/or there are too many oncogene-expressing cells for the immune system to handle, the possibility for exiting senescence provides opportunities for tumour progression. Our findings indicate that deletion of eIF4A2 evokes certain senescence hallmarks, and it is interesting to explore how this might occur. Deletion of eIF4A2 increases ribosome occupancy of mRNAs encoding secretory cargoes and the machinery involved with the processing and transport of these through the ER and Golgi. This, in combination with the tissue expression patterns of eIF4A2 and its association with secretory phenotypes, indicates that a primary function of eIF4A2 is to restrain the secretory translatome; likely to moderate metabolic and proteostatic stress in secretory cells and tissues. Indeed, eIF4A2 deletion increases metabolic activity and key facets of this – increased pyruvate uptake and oxygen consumption – are reversed by rapamycin indicating that this is the energetic cost of an overdriven/de-restrained secretome. Increased entry of pyruvate into the Krebs cycle and the resulting upregulation of oxPHOS is known to contribute BRAF-driven senescence in melanocytes [22], and more recent studies demonstrating that cysteine residues in p21 can act as redox sensors provides mechanistic insight into how one senescence hallmark (increased oxPHOS and ROS generation) can impact another (cell cycle progression) [37]. Furthermore, the unfolded protein response that we observe in eIF4A2 tumours, and in cells derived from these tumours, indicates that de-restraint of secretory mRNAs translation also puts pressure on the protein folding machinery in the ER. Activation of ER-stress sensors (such as PERK, IRE1α and ATF6α) is a characteristic of SASP generation and this, in turn, can drive other senescence hallmarks, including reduced proliferation/cell cycle progression and β-galactosidase expression [38]. The exaggerated secretory behaviour of eIF4A2 knockout cells may, therefore, mimic SASP generation in a way that evokes other senescence behaviours in these cells. Moreover, the SASP has a well-established role in evoking secondary senescence in neighbouring and more distant cells and tissues via paracrine/endocrine mechanisms [39–41]. Indeed, administration of rapamycin to reduce translation of secretory mRNAs in eIF4A2 knockout tumours reduces p21 expression and restores growth. These data are consistent with a paradigm in which overdriven secretory translatomes generate metabolic and proteostatic stresses which then drive other senescence-like behaviours to slow tumour development in vivo.

Inhibition of signalling downstream of mutated KRAS oncogenes is offering possibilities in cancer therapy, but intrinsic and acquired resistance to MAP-kinase inhibition limits use of drugs such as Trametinib as single agents in NSCLC [42]. However, combination of MEK with IGF1R and mTOR inhibition has recently been shown to cause tumour regression in a KRAS-driven pre-clinical mouse model of LuAd [43]. On the face of it, we also show that combined targeting of MEK and mRNA translation opposes KRAS-driven LuAd, but closer analysis hints at a mechanism that is distinct from the ‘one-two punch’ paradigm currently prominent in cancer pharmacology [44]. Crucially, eIF4A2 deletion does not inhibit mRNA translation initiation, but strongly increases synthesis of a secretory translational programme and does so in a manner that evokes senescence-associated behaviours, including metabolic and proteostatic stress. To overcome these stresses, cells in eIF4A2 knockout tumours have re-wired signalling downstream of KRAS by upregulating MAP-kinases and downregulating PI-3 kinase. Thus, it is probable that overdriven mRNA translation in eIF4A2 knockout cells puts them under pressure to minimise activity of PI-3 kinase, thus increasing the dependency on MAP-kinase signalling and rendering eIF4A2 knockout lung tumours exquisitely sensitive to Trametinib. Our observation that human lung cancers with low eIF4A2 also exhibit increased MAP-kinase signalling indicates that eIF4A2 may represent a biomarker for response to therapies targeting MAP-kinases. It will be important in the future to determine the efficacy of MAP-kinase inhibition in targeting LuAd with low eIF4A2 expression, but also other cancers displaying senescence hallmarks linked to increased proteostatic and metabolic stress.

## Materials and Methods

### Assessment of *EIF4A2* mutations across cancer types

cBioPortal for Cancer genomics was used to query for *EIF4A2* mutations across the whole TCGA database [27, 28]. Mutational frequency per cancer type and survival curves were extracted from the cBioportal website. A study with 1053 lung cancer patients from Pan Cancer Atlas was used to query for correlations in mRNA expression of *EIF4A2*, *PI3K* and *MAPK3* genes.

### NSCLC Patient cohort and TMA construction

The cohort (LATTICe-A: Leicester Archival Thoracic Tumour Investigatory Cohort – Adenocarcinoma) used for this study comprises a 994-patient retrospective cohort of resected lung adenocarcinomas and has been previously described [29]. The LATTICe-A cohort was originally constructed under REC 14/EM/1159 (East Midlands REC), and the ongoing management of the resource by the Greater Glasgow and Clyde Biorepository was approved under an amendment granted by the Leicester South REC. Ongoing use of the collection is now managed under REC 16/WS/0207. The full LATTICe-A cohort tissue microarray consisted of 966 tumors from the 1025 tumor cohort. 3 × 1 mm cores were obtained from each case. The entire cohort is represented in 23 tissue microarrays. All tissue microarrays were constructed in quadruplicate using a semiautomated TMArrayer™ (Pathology Devices). Sections were stained with H&E, and all cores were assigned a growth pattern as per The International Association for the Study of Lung Cancer (IASLC) recommendations [45].

### Multiplex imaging

A multiplex immunofluorescence panel consisting of 8 markers was applied to 23 FFPE lung adenocarcinoma TMAs of 4um thickness on TOMO slides (Matsunami TOM-1190). All antibodies were optimised in DAB on the Ventana Discovery Ultra (Roche Tissue Diagnostics, RUO Discovery Universal V21.00.0019) using human lung adenocarcinoma control tissue. Following DAB optimisation, each marker was assigned to an Akoya Opal fluorophore and optimised as a single fluorescent stain to obtain the optimal concentration of fluorophore by checking the exposure time using the Akoya Polaris V1.0.13, with recommended exposure times between 10 and 50ms. Each marker was then assessed in protocol positions 1-8 with its assigned fluorophore to find the optimal position within the multiplex protocol. The finalised multiplex protocol is as follows:

Discovery CC1 (Roche Tissue Diagnostics, 950-123) was applied to the sections for 32 minutes at 95°C for antigen retrieval followed by anti-EIF4A1 (Abcam, ab31217) 1:50, followed by Discovery UltraMap anti-Rb HRP (RUO) (Roche Tissue Diagnostics, 760-4315), detected by Opal 650 (Akoya Biosciences, FP1496001KT) at 1:400, Phospho-p44/42 MAPK (Erk1/2) (Thr202/Tyr204) (D13.14.4E) XP® (Cell Signaling, 4370) 1:100, followed by Discovery UltraMap anti-Rb HRP (RUO) (Roche Tissue Diagnostics, 760-4315), detected by Opal 480 (Akoya Biosciences, FP1500001KT) at 1:50, Anti-EIF4A2 (Abcam, ab31218) 1:100, followed by Discovery UltraMap anti-Rb HRP (RUO) (Roche Tissue Diagnostics, 760-4315), detected by Opal 520 (Akoya Biosciences, FP1487001KT) at 1:300, Anti-AKT1 (phospho S473) (Abcam, ab18206) 1:25, followed by Discovery OmniMap anti-Rb HRP (Roche Tissue Diagnostics, 760-4311), detected by Opal 540 (Akoya Biosciences, FP1494001KT) at 1:100, Anti-eIF4E (phospho S209) [EP2151Y] (Abcam, ab76256) 1:50, followed by Discovery OmniMap anti-Rb HRP (Roche Tissue Diagnostics, 760-4311), detected by Opal 690 (Akoya Biosciences, FP1497001KT) at 1:100, Pan-Cytokeratin (AE1/AE3) (Leica Biosystems, AE1/AE3-601-L-CE) 1:250, followed by Discovery OmniMap anti-Ms HRP (RUO) (Roche Tissue Diagnostics, 05269652001), detected by Opal 620 (Akoya Biosciences, FP1495001KT) at 1:100, Anti-eIF4E antibody [Y449] (Abcam, ab33768) 1:100, followed by Discovery OmniMap anti-Rb HRP (Roche Tissue Diagnostics, 760-4311), detected by Opal 570 (Akoya Biosciences, FP1488001KT) at 1:100, and finally Ki67 (30-9, Ready To Use) (Roche Tissue Diagnostics, 790-4286), followed by Discovery OmniMap anti-Rb HRP (Roche Tissue Diagnostics, 760-4311), detected by Opal 780 (Akoya Biosciences, FP1501001KT) at 1:25. QD DAPI (Roche Tissue Diagnostics, 05268826001, Ready To Use) was applied as a nuclear counterstain. A negative control was applied to a control slide by replacing the primary antibodies above with CONFIRM Negative Control Rabbit Ig, (Roche Tissue Diagnostics, 760-1029) and Negative Control Mouse Monoclonal Antibody (MOPC-211) (Roche Tissue Diagnostics, 760-2014).

Whole slides images were collected at 10x magnification with 7 different filter cubes using the Akoya Bioscience PhenoImager HT, a TMA map was applied to each whole slide scan using Phenochart software (v 2.2.0) and multispectral core images (MSI) were collected at 20x magnification. The MSI were spectrally unmixed using InForm (Akoya Biosciences, v 4.2) and the component image files were imported to Visiopharm (v2021.09.2.10918).

A Deep learning algorithm was trained 288 thousand times on a set of 106 images with diverse tissue morphology. DAPI, autofluorescence and cytokeratin channels were used to train the algorithm to detect regions of necrosis, tumour, stroma, and background tissue categories for subsequent cell level analysis. A deep learning algorithm was trained to detect nuclei and background features on 9 tissue cores with a range of cell types and sizes. The algorithm was trained for 1 million iterations using DAPI and Ki67 features. A separate calculations APP was created to measure area of tumour, stroma, necrosis and the mean intensities of each marker at the cell and regional level. Per-core mean pixel intensities in tumour and stroma were quantified using Visiopharm. Data analysis was conducted in R (version 4.4.2) using the tidyverse package suite (version 2.0.0).

### Mice

All animal experiments were performed in accordance with the UK Animal (Scientific Procedures) Act 1986 and EU direction 2010 and the UK Home Office licences 70/7950, PE47BC0BF and PP7768309. All procedures were ethically reviewed by the animal welfare and ethical review board, University of Glasgow and ARRIVE guidelines were followed for the reporting of in vivo experiments. Animals were housed under controlled conditions (19-22°C room temperature, 45-65% humidity, pathogen free, 12h light/dark cycle) with access to food and water ad libitum. All animals received environmental enrichments including gnawing sticks, plastic tunnels, and nesting material. To reduce pain, suffering and distress, single use needles were used when administering virus and drug treatments, and non-adverse handling techniques were adhered to. All mice used were bred in-house and were of a mixed background. Genotyping was performed by Transnetyx Inc. using ear notches taken for identification purposes at weaning (approximately 3 weeks of age).

The following transgenic alleles were used throughout this study: *Kras^G12D^* (*B6*.*129S4*-*Kras^tm4Tyj^*^/^*^J^*); *Rosa26*LSL-MYC (Gt(ROSA)26Sor^tm1(MYC)Djmy/J^) [11], *Hprt*-lsl-IRFP [17]; *Eif4a1^f^*^l/fl^ and *Eif4a2^f^*^l/fl^ (generated in-house by the CRUK Scotland Institute transgenic/GEMM production service)[6, 10]. For induction of the transgenic alleles, mice between 12 and 16 weeks of age weighing a minimum of 20g were injected with 1 × 10^8^ viral pfu/mouse of the recombinant adenovirus Ad5mSPC-Cre (VVC-Berns-1168). Virus was diluted in 45 μL of MEM and injected intranasally using the calcium phosphate precipitation method [46]. Mice were sampled at specific time points (see figure legends for details) or at a defined clinical endpoint. Lungs were fixed in neutral buffered saline containing 10% formaldehyde through intratracheal injection and posterior immersion in the fixative for 24h. Sample sizes for survival analysis were estimated due to the absence of previous data. Lehr’s quick formula was used for sample estimation at 80% power {n=16(c.v.)2(ln[r.m.])2, assuming a ratio of the means of 0.2 and a coefficient of variation of 0.3. For all other analyses, sample sizes were determined in accordance with the 3Rs and the number of biological replicates was ≥ 3 mice per cohort for all experiments (see Figure legends for details). The Log-rank Mantel-Cox test was used to determine statistical significance on mice survival. Both males and females were used in all experiments. For drug treatments, mice were randomly divided into treatment and control groups and treated by facility staff without knowledge of anticipated outcomes.

For rapamycin treatment, rapamycin (10 mg/kg) diluted in PBS with 5% ethanol, 5% PEG400 (Sigma) 5% Tween 20 (Sigma) or vehicle control was administered via daily *i.p.* injection and tissues were harvested at specific end points (see figure legends for details). For Trametinib treatment, Trametinib (1 mg/kg) diluted in 0.9% NaCl or vehicle control was administered via single daily *i.p.* injection.

### Pearl imaging

All quantifications were performed with Image Studio v5 (LI-COR). Prior to imaging *in vivo*, mice were treated with depilatory cream (Nair) to reduce scattering and absorption of light, as well as to reduce influence of coat colour on individual images. Li-COR supplied filter sets and light sources were used for the PEARL imaging systems. For excitation of iRFP, emission wavelength of 685 nm (700 channel) was used. The detection maxima was 730 nm and had a peak power rating of 0.5 Watt.

### Spatial transcriptomics

Murine liver sections from 3 biological replicates per genotype plus healthy controls were analysed using the digital spatial profiling procedure[47]. Briefly, formalin fixed paraffin embedded tissue sections were treated with 0.1μg/ml proteinase K (AM2546, Thermo Fisher Scientic) followed by heat mediated epitope retrieval and incubated overnight with RNA oligo probes (mouse Whole Transcriptome atlas, Nanostring, GeoMx NGS RNA WTA Mm). Morphological markers were then detected by IF using the conjugated antibodies Alexa532-PanCK, Alexa594-CD45 and the nuclear stain (SYTO13). Slides were imaged at ×20 magnification using the GeoMx digital spatial profiler (DSP) with the integrated software suite. To select regions of interest for eIF4A1 and eIF4A2 expression levels, serial sections were stained and imaged in the Akoya multiplex system (see multiplex methods) using the antibodies Alexa532-PanCK (Nanostring), eIF4A1 (ab31217 abcam), eIF4A2 (ab31218 abcam) and the DNA marker SYTO13 (Nanostring) and imaged at x20 magnification in the GeoMx DSP. Both sections were overlapped and aligned in GeoMx integrated software using PanCK and SYTO13 markers for the alignment. Images in the hybridised slides were then used to select regions-of-interest (ROIs) of 2,500-300,000 μm2 on which the instrument focuses UV light (385 nm), to cleave the UV sensitive probes with the subsequent release of the hybridised barcodes. 37 ROIs per condition (eIF4A2^wt/wt^ and eIF4A2^fl/fl^) and 9 for healthy controls were selected for UV-mediated cleavage and probe collection. Libraries were prepared using GeoMx Seq Code primers (NanoString) and 1× PCR Master Mix (NanoString) and AMPure XP purication. Library quality was checked using an Agilent Bioanalyzer. The libraries were run on an Illumina NextSeq2000 sequencing system at the University of Glasgow Shared Research Facility using a P3 50 cycle kit (Illumina). The FASTQ files from sequenced samples were converted into Digital Count Conversion (DCC) files using the GeoMx NGS pipeline on NanoString’s DND platform. The DCC files were uploaded onto the GeoMx DSP analysis suite (NanoString), where they underwent quality control, filtering, and Q3 normalization. Normalised GeoMx data were downloaded into RStudio (v2023.09.1+494).. Heatmaps were generated using *ComplexHeatMap*. Gene set enrichment analysis (GSEA) was performed using the *fgsea* package using Hallmark curated gene sets.

### Tissue staining and quantification

*Immunohistochemistry (IHC):* All Haematoxylin & Eosin (H&E) and immunohistochemistry (IHC) staining was performed on 4µm formalin fixed paraffin embedded sections (FFPE) which had previously been placed at 60⁰C for 2 hours. The following antibodies were stained on an Agilent AutostainerLink48, eIF4A1 (ab31217, Abcam), eIF4A2 (ab31218, Abcam) and phospho-p44/42 MAPK (pERK) (9101, Cell Signaling). Sections were loaded into an Agilent pre-treatment module to be dewaxed and undergo heat induced epitope retrieval (HIER) using high pH target retrieval solution (High pH TRS K8004, Agilent). All sections were heated to 97⁰C for 20 minutes in the appropriate TRS. After HIER all sections were rinsed in flex wash buffer (K8007, Agilent) prior to being loaded onto the autostainer. The sections underwent peroxidase blocking (S2023, Agilent) for 5 minutes and washed with flex buffer. The specific primary antibody was applied at a previously optimised dilution for 35 minutes (eIF4A1, 1/2000; eIF4A2, 1/1000; pERK, 1/400). The sections were washed with flex wash buffer before application of rabbit envision secondary antibody (K4003, Agilent) for 30 minutes. Sections were rinsed with flex wash then had liquid DAB (K3468, Agilent) applied for 10 minutes. The sections were then washed in water and counterstained with haematoxylin z (RBA-4201-00A, CellPath). Staining for p21(ab107099) took place on a Leica Bond Rx autostainer. All FFPE sections underwent on-board dewaxing (AR9222, Leica) and epitope retrieval using ER2 solution (AR9640, Leica Biosystems) for 20 minutes at 95°C. Sections were rinsed with Leica wash buffer (AR9590, Leica) before peroxidase block was performed using an Intense R kit (DS9263, Leica). Sections for p21 had blocking solution applied from the Rat ImmPRESS kit (MP-7404, Vector Labs) for 20 minutes. Sections were rinsed with wash buffer and then p21 antibody applied at 1/250 dilution. The sections were rinsed with wash buffer and Rat ImmPress secondary antibody applied for 30 minutes. The sections were rinsed with wash buffer, visualised using DAB and then counterstained with haematoxylin in the Intense R kit. H&E staining was performed on a Leica autostainer (ST5020). Sections were dewaxed in xylene, taken through graded ethanol solutions and stained with Haem Z (RBA-4201-00A, CellPath) for 13 mins. Sections were washed in tap water, differentiated in 1% acid alcohol, washed in tap water and the nuclei stained blue with Scotts tap water substitute (in-house). After washing with tap water sections were placed in Putt’s Eosin (RRSP34-F, Atom Scientific) for 3 minutes. To complete H&E and IHC staining sections were rinsed in tap water, dehydrated through graded ethanol’s and placed in xylene. The stained sections were coverslipped in xylene using DPX mountant (SEA-1300-00A, CellPath). Immunofluorescence staining: antigen retrieval was performed using Citrate Buffer (10mM Citric Acid, pH 6, 0.05% Tween) for 20 minutes at 90oC and tissue was blocked in 1% BSA for 1h at RT and incubated with primary antibody overnight at 4oC. Tissue was incubated with secondary antibodies (Alexa-488 or Alexa-647, Thermo Fisher, 1:400) before being mounted with VECTASHIELD containing DAPI (Vector Laboratories). Antibodies used were: eIF4A2 (abcam 31218, 1:100), KRasG12D (#14429 cell signalling, 1:50) and p21 (abcam 107099, 1:100). High-content automated microscopy: Stained slides were imaged at ×20 magnification using the Opera Phenix High-Content Screening System (Harmony High-content Imaging and analysis software version 4.9, PerkinElmer) and images were quantified using Columbus Image Data Storage and Analysis System (PerkinElmer version 2.8.0). IHC scanning and analysis: Stained slides were scanned using the Leica Aperio AT2 slide scanner and analysed using HALO® image analysis software. Tumour burden was quantified in H&E-stained sections using HALO® Area quantification v2.4.2. p21 protein expression levels were quantified using HALO® cytonuclear v2.0.9 analysis trained to quantify the optical density of nuclear (p21) staining.

### Cells lines, antibodies and reagents

*Mouse cell lines:* KM cells were isolated from *Kras*^LSL-G12D/WT^; *Rosa26*^LSL-MYC/LSL-MYC^ mouse tumours with either *Eif4a2*^wt/wt^ or *Eif4a2*^fl/fl^ allele. One clinical endpoint male mouse was used to create each cell line (KM^NTC/4A2cr^; KM^A2+/+^; KM^A2−/−^). Briefly, left lobes from clinical endpoint mice were mechanically disrupted and passed through a 70um mesh. Cells were then resuspended in DMEM supplemented with 10% fetal bovine serum (FBS), 2 mM glutamine, 100 IU/ml penicillin and 100 μg/ml streptomycin and cultured in the presence of cholera toxin to minimise fibroblast growth. Further enrichment for epithelial cells vs fibroblasts was obtained using differential trypsinization. Cell identity was assessed by expression of KRAS^G12D^, SPC and overexpression of MYC against a breast cancer mouse cell line. Expression of the iRFP reporter was also confirmed in the KM cells. All cell lines were tested for the presence of mycoplasma.

For Western blots, the following antibodies were used at a dilution of 1:1000: eIF4A1 (abcam 31217), eIF4A2 (abcam 31218), pmTORC1 (Cell signalling 2971), pPERK (cell signalling 3179), ATF4 (Cell signalling 11815), pAKT (Cell signalling 3787), pERK1/2 (Cell signalling 9106), ERK1/2 (Sigma M5670) GAPDH (Sigma G8795), β-actin (Sigma A1978), p21 (Abcam ab107099), p16 (Abcam ab211542)

#### CRISPR approaches

Guide RNA (gRNA) sequences targeting *Eif4a2* were cloned into a lentiCRISPR vector (Addgene 52961). gRNAs were as follows: eIF4A2 (forward) CACCGGAAGCCCCTCACATTGTTGT; eIF4A2 (reverse) AAACACAACAATGTGAGGGGCTTCC. To produce lentivirus encoding Cas9, along with the desired gRNAs, HEK 293FT cells were used as the host packaging cell line. Virus-containing HEK 293FT supernatant was then filtered through a 0.45μm PTFE filter membrane, polybrene supplemented to a final concentration of 4μg/ml and the medium incubated with recipient KM cells overnight at 37°C/ 5% CO2. Virus transduced cells were then selected using puromycin (2μg/ml), the antibiotic selection medium being replenished every 2 days, and the cells were passed according to their confluence to generate stable cell lines. To ensure lack of eIF4A2 expression, stable cell lines were clonally selected by plating individual cells in 96-well plates, allowing growth and subsequent passaging to 24 and 6-well plates and assessing eIF4A2 expression by western blot. To eliminate clonal effect, 4 clones negative for eIF4A2 were pooled to form the eIF4A2-crispr LKM cell line.

#### Cell growth, treatments and ROS measurements

Cells were seeded at 1×10^4^ cells per well in 24-well culture plates and incubated at 37°C/ 5% CO2 in a humidified incubator for 12 hours to adhere. Cells were treated with either Rapamycin 1uM, eft226 100nM, Trametinib 100nM or vehicle control and labelled with the ROS marker CellROX green (Invitrogen) at 2.5μM and then transferred to an IncuCyte ZOOM live-cell imaging system (Sartorius). Four technical replicates were analysed per condition. Images were acquired (10x objective) every hour over a period of 72h and analysed using the Incucyte Base Analysis Software (Version 2023A Rev2), Confluence was calculated as a % of the phase image area covered by cells and ROS as the Integrated density of Cell Rox signal (fluorescent units/μm2). For β-galactosidase activity measurements, the Cell Event Senescence Green Detection kit (Invitrogen) was used following manufacturer’s instructions.

### Spheroids

To determine cells’ growth capacity, 1,000 KM^NTC^, and KM^4A2cr^ cells were suspended in DMEM F12 Advance (Thermo Fisher Scientific) supplemented with 25 ng/ml B-27 (Thermo Fisher Scientific), and 1X N-2 (Gibco) and seeded into 2% agarose-coated 12-well plates. After incubation at 37°C for 12 days, spheres >70 μm diameter were scored.

### Colony formation

KM^NTC^, and KM^4A2cr^ cells were seeded at a density of 300 cells per well in 6-well plates with three technical repeats per experiment and left to adhere overnight. Cells were left to form colonies for 1 to 2 weeks prior to methanol fixation and crystal violet staining. Visible colonies consisting of a minimum of 50 cells were counted manually.

### Ribosome footprinting

KM^NTC^, and KM^4A2cr^ cells were plated in 10cm plates and allow to reach 90% confluency. Cells were then gently lifted, pelleted in PBS and lysed in 500 µl ice-cold lysis buffer (15 mM Tris-HCl pH 7.5, 15 mM MgCl2, 150 mM NaCl, 1% Triton X-100, 0.05% Tween 20, 2% n-Dodecyl β Maltoside (Thermo Fisher Scientific, 89903), 0.5 mM DTT, 100 µg/ml cycloheximide, 1× cOmplete EDTA-free Protease Inhibitor Cocktail (Merck, 11836170001), 200 U/ml RiboLock RNase Inhibitor (Thermo Fisher Scientific, EO0381)) on ice for 5 min with agitation every minute. Lysates were centrifuged at 4°C for 5 min at 12,000 g and 500 µl supernatant was pipetted into a fresh 1.5 ml tube. For cytoplasmic RNA samples, RNA was extracted from 25 µl of lysate with 975 µl TRIzol, as per the manufacturer’s instructions. A 20-µl aliquot of lysate was also added to 5 µl of 5× SDS gel-loading buffer for western blotting to check for depletion of p. The remaining lysate was digested with 5 µl Ambion RNase I (cloned, 100 U/µl; Thermo Fisher Scientific, AM2295) at 22°C for 15 min, with agitation at 600 rpm in a thermo-mixer. The digestion was stopped with 10 µl SUPERaseIn RNase Inhibitor (20 U/μl; Thermo Fisher Scientific, AM2696) and samples were loaded onto a 10–50% sucrose gradient, containing 15 mM Tris-HCl pH 7.5, 15 mM MgCl2, 150 mM NaCl, and 100 µg/ml cycloheximide, prepared using a BioComp gradient station and cooled to 4°C for at least 1 h prior to use. Samples were centrifuged in a Beckman XPN-90 Ultracentrifuge with an SW40Ti rotor at 38,000 rpm for 2 h at 4°C. Fractions of 0.5 ml were collected with a Biocomp gradient station and Gilson FC 203B fraction collector. Fractions containing the 80S ribosome peak were extracted with acid phenol:chloroform, followed by two chloroform washes. RNA was precipitated with 2 µl glycogen (Roche 10901393001), 300 mM NaOAc pH 5.2, and an equal volume of isopropanol overnight at −20 °C.

RNA was pelleted by centrifugation at 12,000 g for 45 min at 4 °C. The supernatant was removed with a pipette, and the pellet was washed twice with 70% ethanol and dissolved in 10 µl RNase-free water. RNA was diluted with 10 µl 2× urea buffer (Thermo Fisher Scientific, LC6876), heated at 80°C for 90 s, placed immediately on ice, and then loaded onto a pre-run 15% TBE-urea gel (Thermo Fisher Scientific, EC68852BOX) and ran at 200 V for 1 h alongside custom 28 nt (AGCGUGUACUCCGAAGAGGAUCCAACGU) and 34 nt (GCAUUAACGCGAACUCGGCCUACAAUAGUGACGU) RNA markers. The gel was stained with 1× SYBR gold (Thermo Fisher Scientific, S11494) and imaged on a Typhoon FLA 7000. An image was printed to size to allow bands, inclusive of 28 nt and exclusive of 34 nt markers, to be cut from the gel, placed into a 1.5 ml RNA low-binding microcentrifuge tube, and crushed with a scalpel. RNA was eluted from the crushed gel pieces in 500 µl extraction buffer (300 mM NaOAc pH 5.2, 1 mM EDTA, 0.25% SDS) overnight at 16°C at 600 rpm in a thermo-mixer. Gel pieces were filtered out with a Costar Spin-X centrifuge tube filter (0.45 µm; Scientific Laboratory Supplies Ltd, 8163) and RNA was precipitated with 2 µl glycogen and 500 µl isopropanol overnight at −20 °C.

Precipitated RNA was pelleted, washed, and dissolved in 13.5 µl RNase-free water, as above. To deplete rRNA the following was added to the 13.5 µl RNA; 5 µl hybridisation buffer (10mM Tris-HCl pH7.5, 1mM EDTA, 2M NaCl), 1 µl RNasin Plus Ribonuclease Inhibitor (Promega N2615) and 0.5 µl biotinylated DNA oligo pool (100 uM total DNA with rRNA_depl_1 and rRNA_depl_2 at a 3:1 molar ratio compared to all other oligos), that had been denatured at 95 °C for 3min and then placed immediately on ice. The mix was incubated for 10min at 68 °C at 1250 rpm in a thermo-mixer and then allowed to cool slowly to room temp by turning off the thermo-mixer. rRNA was depleted with 160 µl Dynabeads MyOne Streptavidin C1 (Thermo Scientific, 65001) as per manufactures’ instructions. The depleted non-RNA was precipitated with 2 µl glycogen and three times the volume of ethanol.

Precipitated RNA was again pelleted, washed, and dissolved in 43 µl RNase-free water, as above. RNA was heated at 80°C for 90 s and immediately placed on ice before undergoing 5′ phosphorylation and 3′ dephosphorylation with 1 µl T4 PNK (NEB, M0201S), 5 µl 10× PNK buffer, and 1 µl SUPERaseIn RNase Inhibitor (20 U/μl) at 37°C for 35 min, with 5 µl 10 mM ATP added for the final 20 min. RNA was extracted with acid phenol:chloroform and isopropanol-precipitated as above.

Purified RNA was input into the Bioscientific Nextflex small RNA v3 kit (NOVA-5132-06), using the alternative step F bead clean-up, 15 PCR cycles, and gel-extraction option.

Cytoplasmic RNA samples were run on an Agilent TapeStation to check RNA integrity. RNA concentration and sample purity were measured on a NanoDrop spectrophotometer. rRNA was depleted from 1 µg cytoplasmic RNA with the RiboCop v2, and sequencing libraries were prepared from the rRNA-depleted RNA with the Corall Total RNA-Seq Library Prep Kit v1 (Lexogen, 095.96) using 13 PCR cycles.

Final cytoplasmic RNA and RPF libraries were quantified on an Agilent TapeStation, with a High Sensitivity D1000 tape (Agilent), and sequenced single-end on an Illumina NextSeq500 instrument with a 75 cycles high-output kit.

### Bioinformatic processing of ribosome footprinting data

Ribosome-protected fragments (RPF) and total cytoplasmic RNA files were processed using the https://github.com/Bushell-lab/Ribo-seq GitHub package, using R version 4.3.1. Briefly, fastQC was used to QC-check raw fastq files. Cutadapt (version 1.18) was used to remove adaptors, trim 3’ bases with Phred scores < 20, and discard reads fewer than 30 and more than 50 bases after trimming. UMI-tools (version 1.0.1) [48]was used to extract unique molecular indexes (UMIs) (4 nt of random sequence at the start and end of every RPF read) from the reads and appended to the read name. Reads were aligned with BBmap (version 38.18), first to remove reads that align to either rRNA or tRNA sequences. Non-rRNA/tRNA reads were then aligned to a filtered protein-coding FASTA (see below), with only the most abundant transcript per gene (calculated from cytoplasmic RNA samples). Samtools (version 1.9) was used to sort and index the resulting BAM files and UMI tools used to deduplicated with the directional method. The number of reads with the 5’ end at each position of every transcript was then counted. Protein coding aligned read lengths peaked at 29-31 nt and lengths 28-33 were used in this analysis, which showed strong CDS enrichment and periodicity. The offset for each read length was determined to be 12 nt for read lengths 23-30 nt, and 13 nt for 31-33 nt. Total counts across the entire CDS, except the first 20 and last 10 codons, were summed together and used as input into DESeq2 to test for differential expression. This was carried out separately on either the RPF or cytoplasmic RNA samples to calculate log2FCs and plot PCAs. Gene Set EnrichmentAnalysis (GSEA) was carried out using R package fgsea version 1.8.0, and comparison gene sets obtained from available sources (Gene Ontology Consortium). The paired cytoplasmic RNA samples were processed as the RPFs but with the following exceptions: UMIs were 12 nt at the start of each read only. No maximum read length was set when trimming reads with Cutadapt. Reads were aligned to a filtered protein-coding transcriptome (see below) with Bowtie2 (version 2.3.5.1)[49], using the parameters recommended for use with RSEM, which are --sensitive –dpad 0 --gbar 99999999 --mp 1,1 --np 1 --score-min L,0,-0.1. Gene and isoform level quantification was performed using RSEM (version 1.3.3)[50]. The isoform quantification was used to determine the most abundant transcript per gene (see below), but differential expression was measured at the gene level with DESeq2 69. The gencode.vM27.pc_transcripts.fa file was downloaded from https://www.gencodegenes.org/mouse/release_M27.html and filtered to include only transcripts that had been manually annotated by HAVANA and that have a 5’UTR, a 3’UTR, a CDS equally divisible by 3, an nUG start codon, and a stop codon. All PAR_Y transcripts were also removed. The cytoplasmic RNA-seq data was then aligned to this filtered FASTA and the most abundant transcript per gene was determined, based on the mean TPMs across all samples from the RSEM output. The RPF reads were then aligned to a FASTA containing only the most abundant transcript for each gene.

### Oxygen Consumption Rate

Oxygen consumption rates (OCRs) were determined using the Seahorse Extracellular Flux Analyzer (Agilent) according to the manufacturer’s instructions. Briefly LKM cells were plated at 3 × 104 cells/well in a 96-well plate, and assay medium was prepared with phenol-free DMEM. OCR was measured during stepwise injection of 1 µM oligomycin, 1 µM carbonyl cyanide-4-(trifluoromethoxy) phenylhydrazone (FCCP), and 1 µM rotenone/antimycin A (Sigma-Aldrich). After completion, cells were lysed and OCR was normalised to protein content determined using a Pierce bicinchoninic acid protein assay kit (Thermo Fisher Scientific) according to the manufacturer’s instructions. Basal OCRs were calculated by subtracting the average post-rotenone/antimycin A OCR value from the average initial OCR value.

### Metabolomics

*Cell extracts:* 200,000 cells were seeded in 2 mL medium per well in 6-well plates and 24h later the medium was replaced with 7 mL DMEM (Thermo 10313021) supplemented with 2 mM glutamine and 10% FBS. 48h after seeding, cells were washed three times with ice-cold PBS and incubated for 5 min at 4C with 600 µL of LC–MS extraction solution (20% water, 50% methanol and 30% acetonitrile). 3 technical replicates were used per condition in each of 3 independent experiments.

Extracts from extracellular media: 500,000 cells were seeded in 2 mL medium per well in 6-well plates and the medium was replaced 24h later with 2 mL DMEM (Thermo 10313021) supplemented with 2 mM glutamine and 10% FBS. 48h after seeding, 20 µL of the extracellular medium was extracted in 980 µL of LC–MS extraction solution (20% water, 50% methanol and 30% acetonitrile). 6 technical replicates were used per condition in each of 5 independent experiments.LC–MS metabolomics: A Q Exactive Plus Orbitrap Mass Spectrometer (Thermo Fisher Scientific) was employed coupled with an Ultimate 3000 high performance liquid chromatography (HPLC) system (Thermo Fisher Scientific). The samples were injected (5 μL) and separated on a ZIC-pHILIC column (SeQuant; 150 mm by 2.1 mm, 5 μm; Merck 521 KGaA, Darmstadt, Germany) coupled with a ZIC-pHILIC guard column (SeQuant; 20 mm by 2.1 mm) 45°C. The chromatographic separation was performed with a resolution of 35,000 (at 200 m/z) with electrospray ionisation and polarity switching, to detect both positive and negative ions over a mass range of 75-1000 m/z. The metabolites were separated with a 15 min mobile phase gradient, which started at 80% acetonitrile / 20% ammonium carbonate (20 mM, pH 9.2), and decreased to 20% acetonitrile / 80% ammonium carbonate with a flow rate of 200 μL/min (total run time of 24.5 min).

Targeted metabolomics analysis was performed using Skyline (23.1.0.455) [51], and the peak areas of metabolites were determined by using the m/z of the singly charged ions (extracted ion chromatogram, ±5 ppm) and the retention time from our in-house metabolite library. The data was normalised to cell counts and plotted as fold change relative to the vehicle-treated NTC cells from each biological repeat.

## Acknowledgments

We would like to thank core staff in the Biological Services Unit, the Beatson Advanced Imaging Resource (BAIR), Molecular Services, Histology Facility and Central Services (CRUK-Scotland Institute) for their support which facilitated the work described in this manuscript. We would also like to thank the NHSGGC Biorepository, MRC Toxicology Unit, and the University Hospital Leicester for the management of the LATTICe TMA cohort and Madhumita Das, Claire Wilson, and Marco Sereno for the construction of the LATTICe TMA Cohort. This paper was critically reviewed by Catherine Winchester (CRUK Scotland Institute).

## Funding

This work was funded by Cancer Research UK core programme funding to JCN (A18277 and A28291) and the Medical Research Council (MR/P01058X/1). We acknowledge the Cancer Research UK Glasgow Centre (C596/A18076) and the BSU facilities at the Cancer Research UK Beatson Institute (C596/A17196 and A31287). LP, MM, RD, JAW, EJ, LMc, MH, JM, LM, DaS, DoS and MB were funded by Cancer Research UK core programme 31287. BK and DM were funded by CRUK project grant A27603 and the National Mouse Genetics Network Cancer Cluster (MC/PC21042). IP, LOJ, RP and JLQ were funded by Mazumdar-Shaw Chair Fund. CW, HL, and NJ were funded by Cancer Research UK (C55370/A25813)

## Author contributions

Experimental design: LP, JCN, MM, MB, DLM

Investigation: LP, RD, MM, IP, JAW, BK, EJ, LMcG, CW, HL, MH, LM, LOJ, NJ, David S, Douglas S, RP

Reagents: LP, JCN, DLM, JLQ

Funding acquisition: JCN, JLQ, DLM, MB

Writing – original draft: LP & JCN

Writing – review & editing: LP, JCN, MM, RD, DLM, JLQ, MB

## Competing interests

Authors declare that they have no competing interests.

## Data and materials availability

The data supporting the findings of this study are available within the article and its supplementary information files and from the corresponding author upon request.

## Supplementary figures

**Figure S1.**
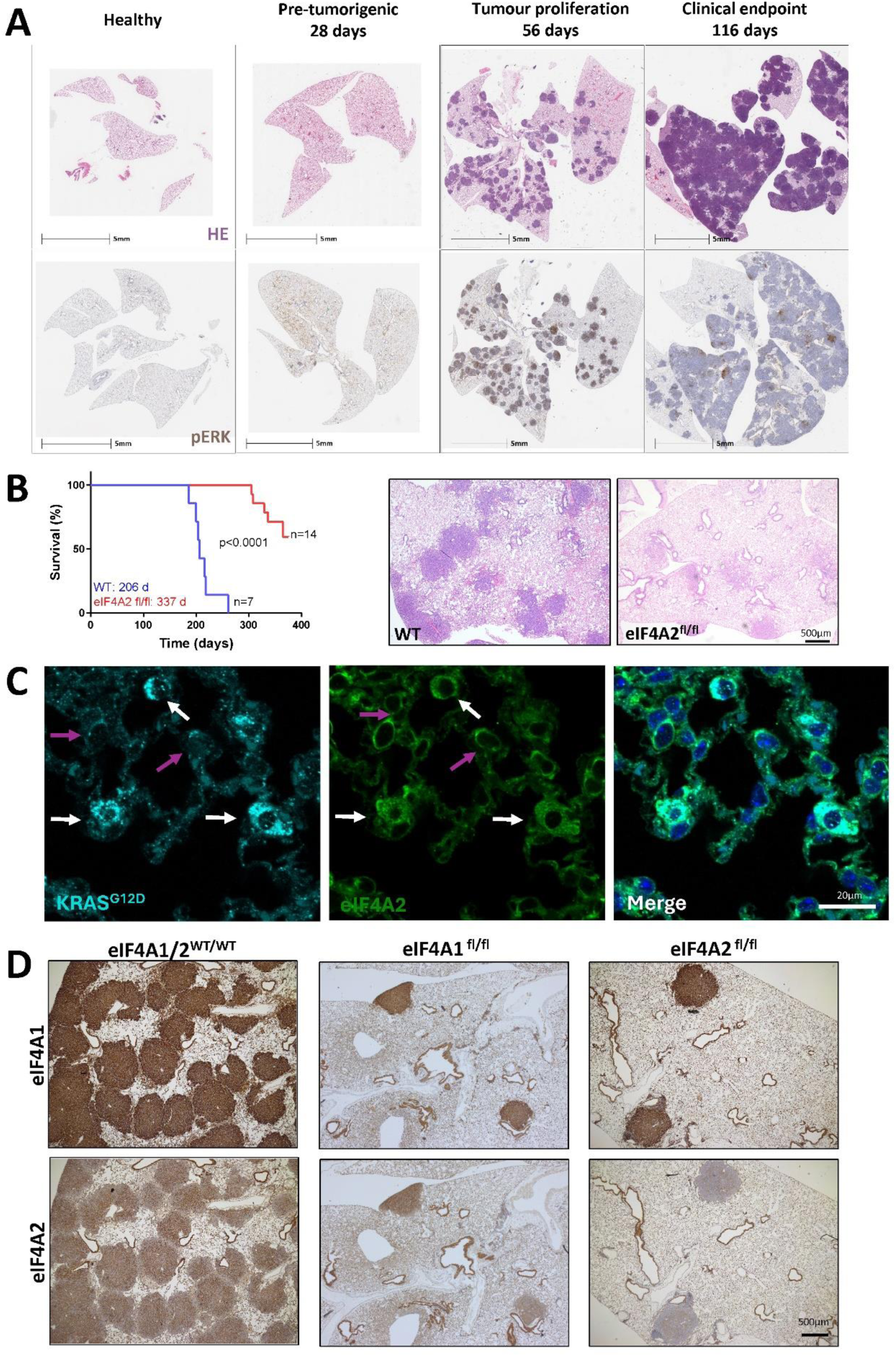
KRAS-driven mouse of lung cancer and influence of eIF4A paralogues. **(A)** Ad5-SPC-CRE was administered intranasally to *KRAS*^LSL-G12D/WT^; *Rosa26*^LSL-MYC/LSL-MYC^ (KM) mice at a titre (1 × 10^8^ plaque forming units (pfu)/mouse) sufficient to evoke recombination in ≈5% alveolar type II cells. Some mice were left uninduced (healthy). Mice were sacrificed at the indicated time points and lungs removed and fixed. Tumour burden was visualised using H&E and phospho-ERK by immunohistochemistry. **(B)** *KRAS*^LSL-G12D/WT^; eIF4A2^WT/WT^ or *KRAS*^LSL-G12D/WT^; eIF4A2^fl/fl^ mice were induced with Ad5-SPC-CRE as for (A). Mice were sacrificed at clinical endpoint or, in the case of several animals from the *KRAS*^LSL-G12D/WT^; eIF4A2^fl/fl^ cohort at 375 days following induction with Ad5-SPC-CRE. The H&E images display the tumour burden of a *KRAS*^LSL-G12D/WT^; eIF4A2^WT/WT^ mouse at clinical endpoint (left panel) and the lung field of a *KRAS*^LSL-G12D/WT^; eIF4A2^fl/fl^ mouse taken 375 days following Ad5-SPC-CRE administration that had not displayed clinical signs of lung cancer. The statistical test for the Kaplan-Maier analysis in is Logrank (Mantel-Cox) and cohort sizes were N=7 individual mice for eIF4A2^wt/wt^ (WT) and N=14 for eIF4A2^fl/fl^. Only 9 of the 14 mice in the eIF4A2^fl/fl^ cohort had reached clinical endpoint by 375 days following Ad5-SPC-CRE administration when the experiment was terminated. **(C)** Ad5-SPC-CRE was administered to KM mice. Mice were sacrificed 28 days following and mutant KRAS^G12D^ (blue) and eIF4A2 (green) were visualised in lung slices using immunofluorescence. eIF4A2 expression is increased in KRAS^G12D^ positive cells (white arrows) and reduced in KRAS^G12D^ negative cells (magenta arrows). **(D)** KM mice that were either eIF4A1/2^WT/WT^, eIF4A1^fl/fl^ or eIF4A2^fl/fl^ were induced with Ad5-SPC-CRE. Mice were sacrificed 56 days following Ad5-SPC-CRE administration and eIF4A1 and eIF4A2 visualised in the lung using immunohistochemistry.

**Figure S2.**
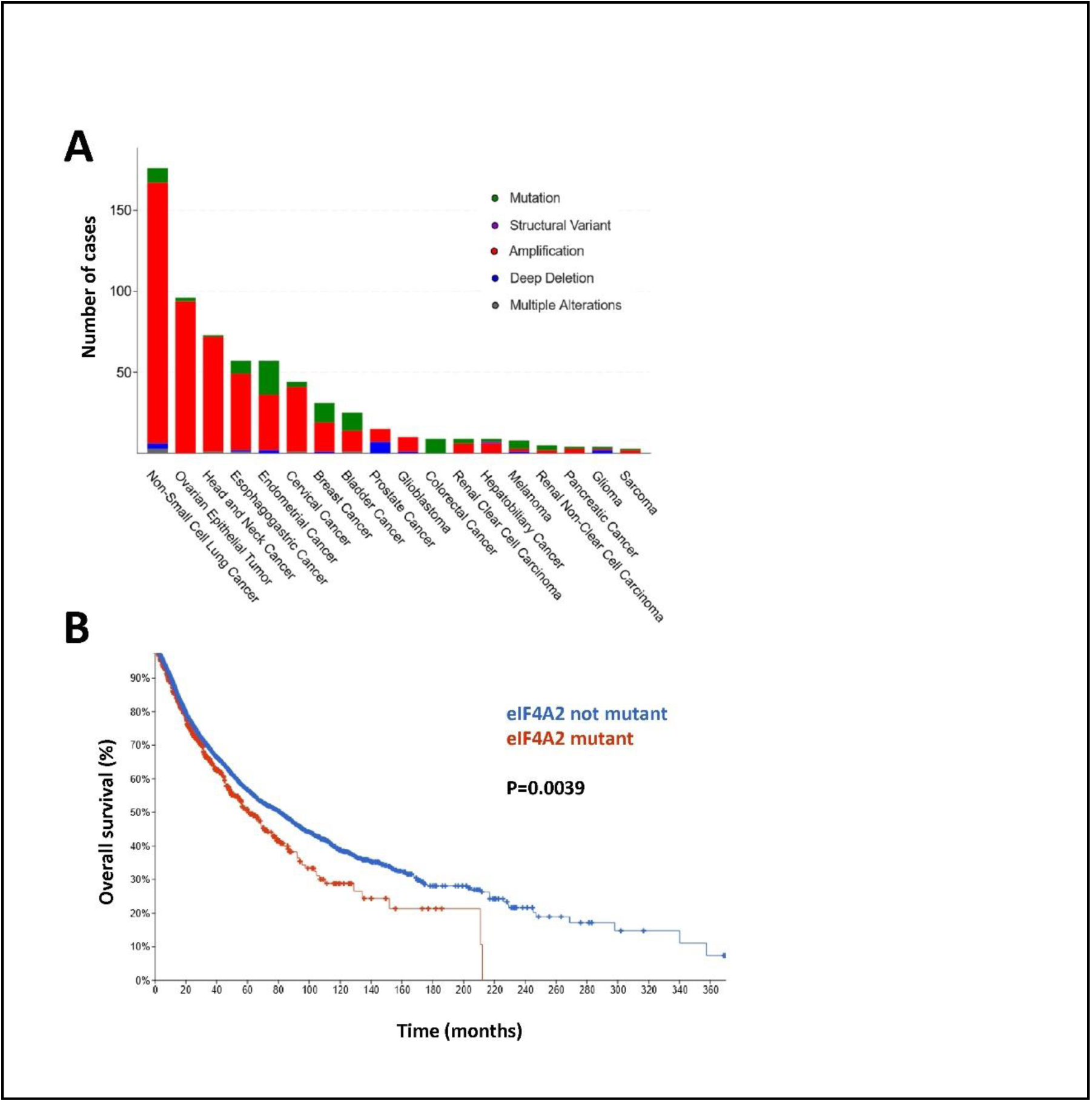
Distribution of eIF4A2 mutations across cancer types. **(A)** Percentages of eIF4A2 mutations were obtained from cBioportal by querying the TCGA dataset (10953 patients) [13]. eIF4A2 mutations are present in 6% of cancers on average, but in 16.7% of Non-Small-Cell Lung Cancer. The majority of eIF4A2 mutations in NSCLC lead to amplification of the gene. **(B)** The TCGA dataset was used to interrogate the overall survival of patients with NSCLC which do (red, n=642) and do not (blue, n=10318) display mutation of the eIF4A2 gene. Patients with NSCLCs harbouring mutations in eIF4A2 exhibit poorer overall survival. The statistical test for the Kaplan-Maier analysis in is Logrank (Mantel-Cox).

**Figure S3.**
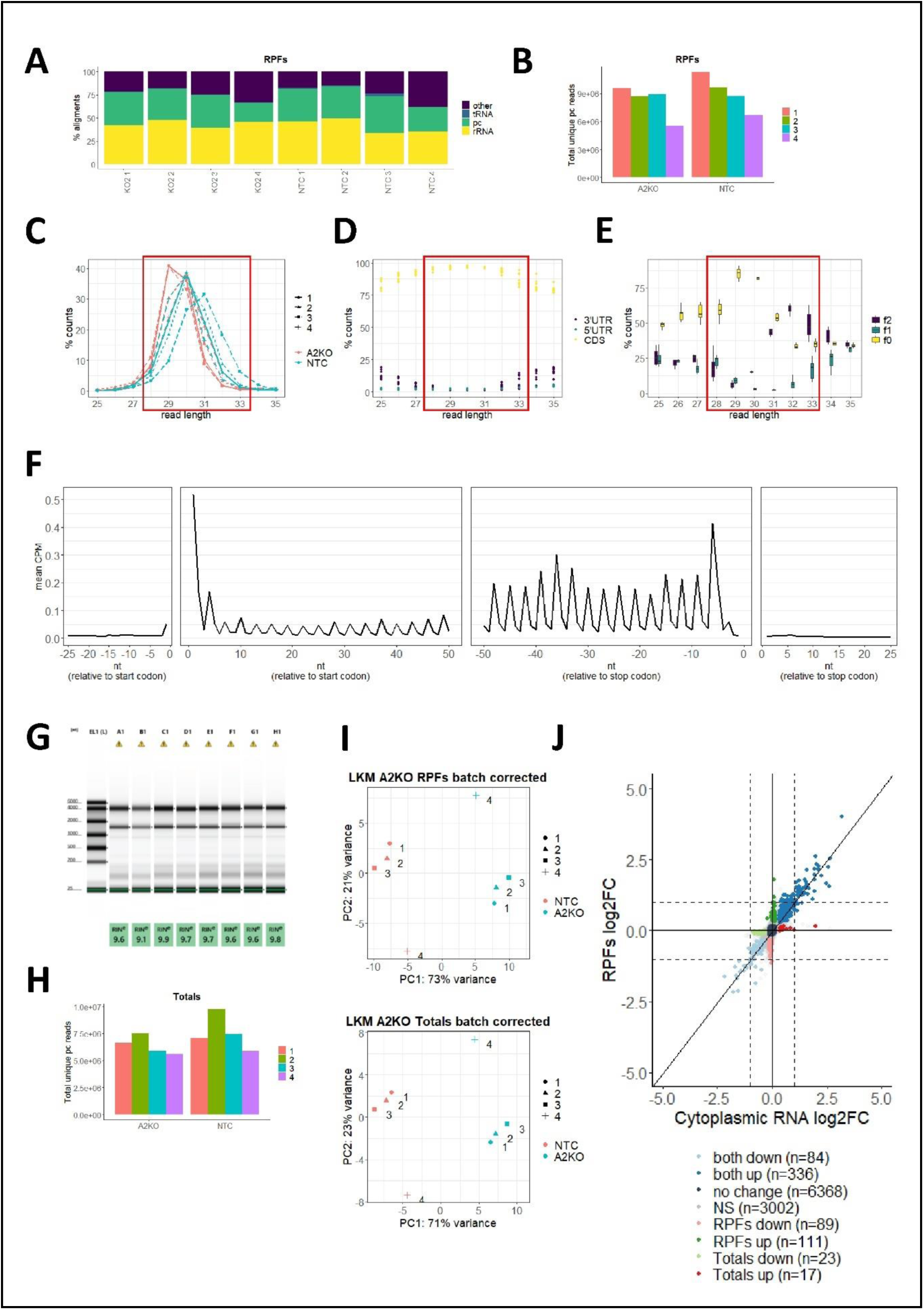
RiboSeq Quality Control. **(A)** Alignment percentages across all ribosome-protected fragment (RPF) samples. **(B)** Total number of unique protein-coding reads for all RPF samples. **(C)** Distribution of read lengths for all protein-coding RPF samples. **(D)** Proportion of reads aligning to the 5’UTR, CDS, or 3’UTR for various read lengths. **(E)** Boxplot showing the percentage of RPF reads within each coding sequence frame, categorized by read length. The red boxes in panels C-E highlight the read lengths selected for further analysis. **(F)** Mean counts per million (CPM) for all transcripts and samples at the indicated positions, after applying the offsets described in the methods section, demonstrating periodicity specifically within the CDS. **(G)** Tapestation file indicating that total cytoplasmic RNA remains intact, with high RNA Integrity Number (RIN) values. **(H)** Total count of unique protein-coding reads from the total cytoplasmic RNA samples. **(I)** Batch corrected PCA plots showing distribution of the replicates per condition in RPF and total RNA. **(J)** KM^4A2CR^/KM^NTC^ Log2FC of RPFs vs total RNA showing increased translational efficiency in KM^4A2CR^.

**Figure S4.**
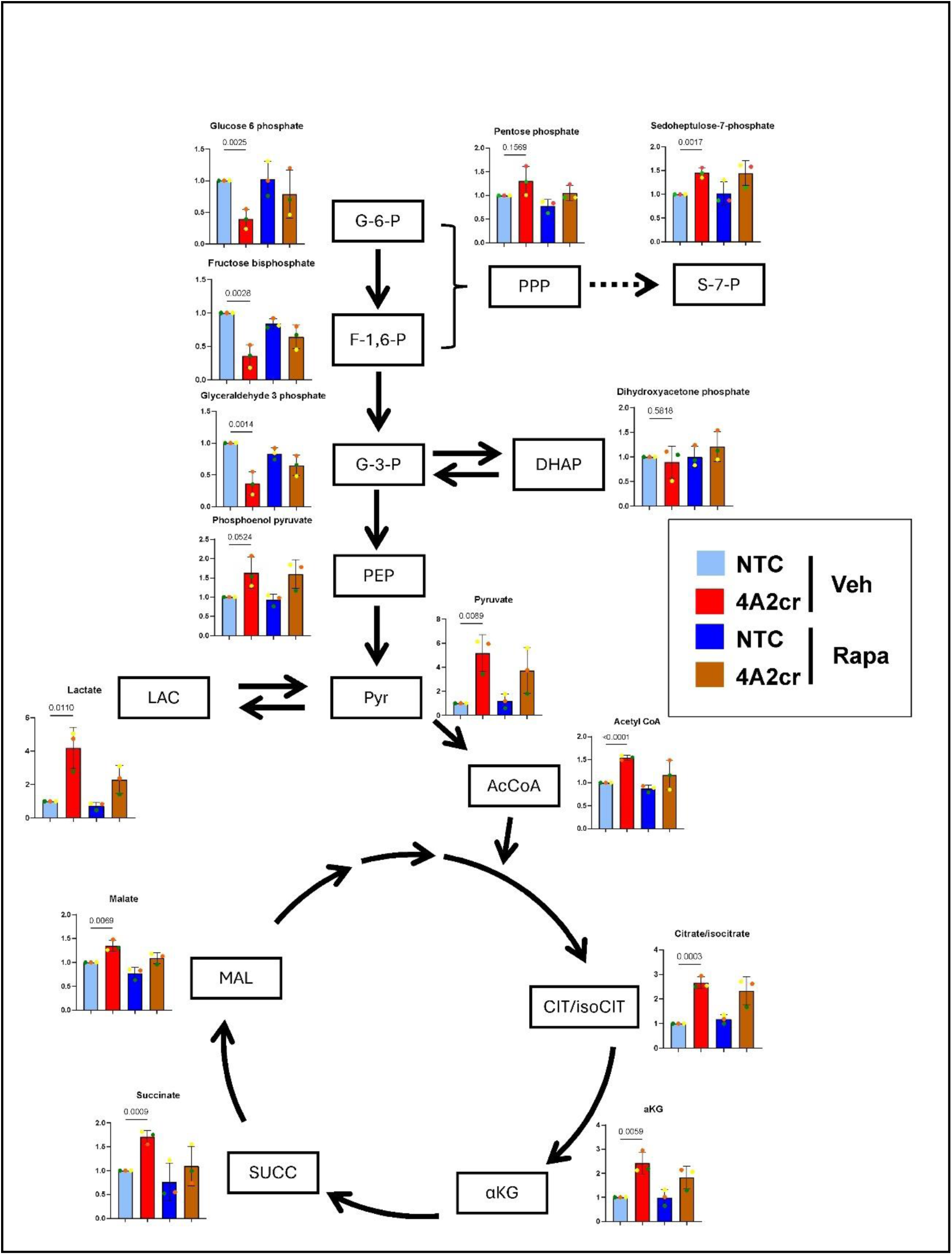
eIF4A2 depletion influences metabolite levels. KM^NTC^ or KM^4A2cr^ cells were plated onto six-well dishes and allowed to adhere overnight. Adherent cells were incubated at 37°C for 24 hr in the presence and absence of rapamycin (1 μM) or vehicle control (Veh.). Cell lysates were used for extraction of polar metabolites. LC-MS was used to determine metabolites and levels of these are plotted as peak areas normalised to those of KM^NTC^ cells. Bars are mean ± SEM, N=3, each coloured dot denotes an individual experiment. Statistical test is ANOVA with Tukey’s multiple comparison.

**Figure S5.**
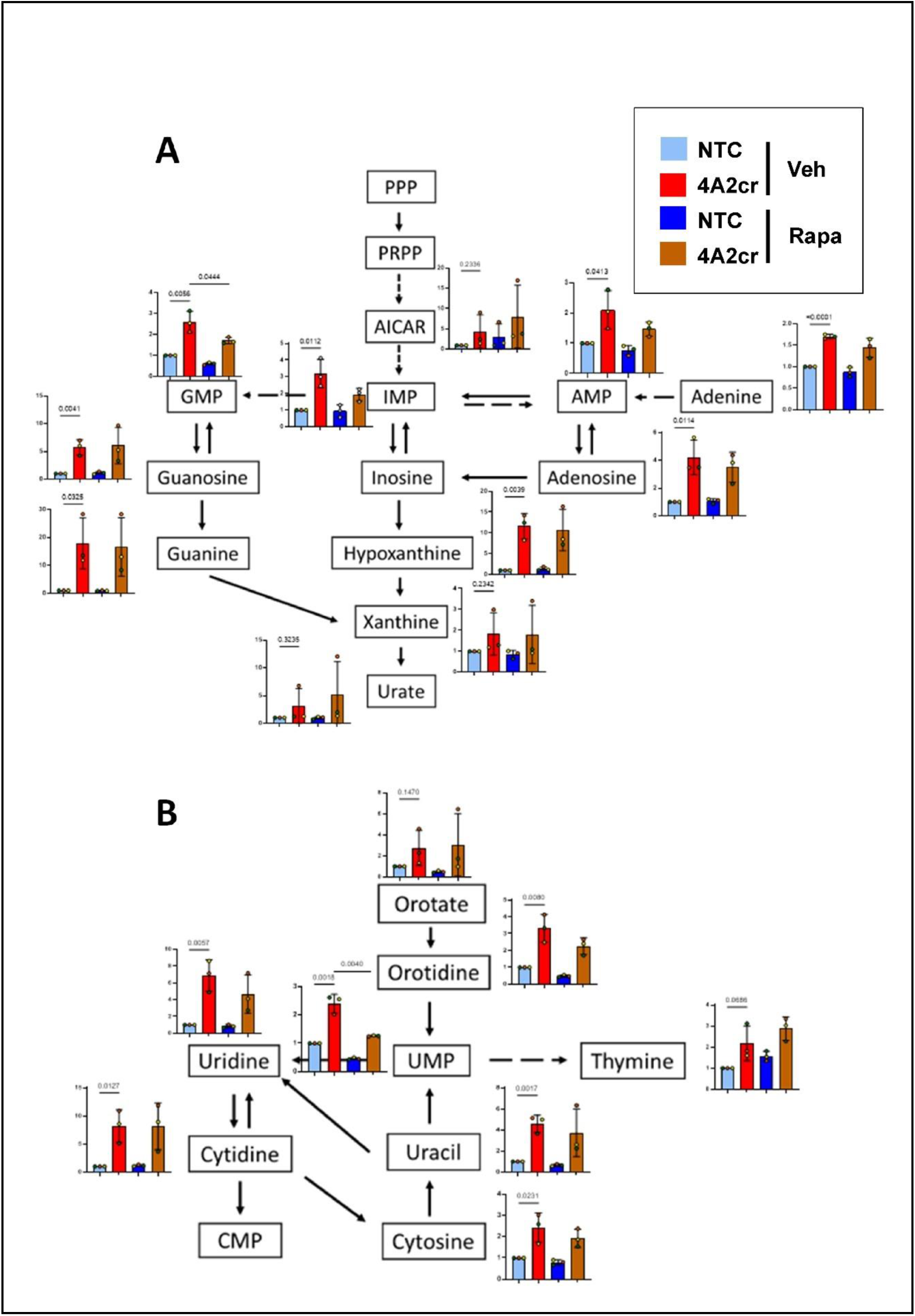
eIF4A2 influences nucleotide synthesis and metabolism. KM^NTC^ or KM^4A2cr^ cells were plated onto six-well dishes and allowed to adhere overnight. Adherent cells were incubated at 37°C for 24 h in the presence and absence of rapamycin (1 μM) or vehicle control (Veh.) and polar metabolites were extracted after treatment. LC-MS was used to determine metabolites known to be involved in purine **(A)** and pyrimidine **(B)** synthesis and/or metabolism and levels of these are plotted as peak areas normalised to those of KM^NTC^ cells. Values are mean ± SEM, N=3, each coloured dot denotes an individual experiment. Statistical test is ANOVA with Tukey’s multiple comparison.

**Fig. S6.**
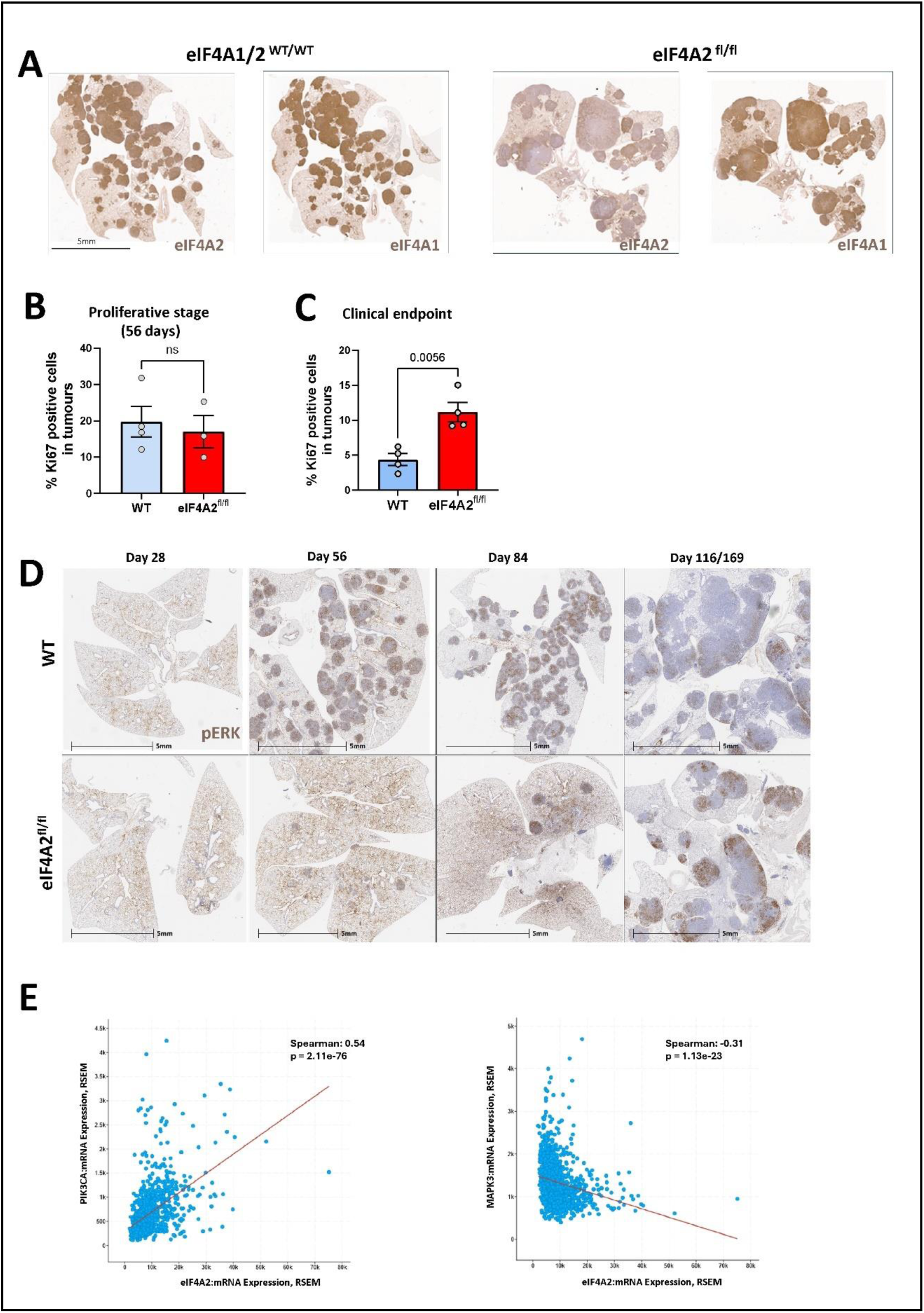
Characteristics of eIF4A2 knockout tumours. **(A-D)** Ad5-SPC-CRE was administered intranasally to *KRAS*^LSL-G12D/WT^; *Rosa26*^LSL-MYC/LSL-MYC^ (KM) mice that were either *eIF4A2*^WT/WT^, or *eIF4A2*^fl/fl^. Mice were sacrificed at clinical endpoint and lungs removed and fixed and stained for eIF4A1, eIF4A2 **(A)** or phospho-ERK **(D)** using immunohistochemistry. For **(B)** and **(C)** mice were sacrificed at 56 days post induction or clinical endpoint and stained for the proliferative marker Ki67. **(E)** A NSCLC study (1053 patients) from Pan Cancer Atlas was queried in c-Bioportal for RNA expression of eIF4A2, PI3K and ERK1 (MAPK3). Graphs show Pearson and Spearman correlations of eIF4A2 with either PI3K (positive correlation) or ERK1 (negative correlation).

**Fig. S7.**
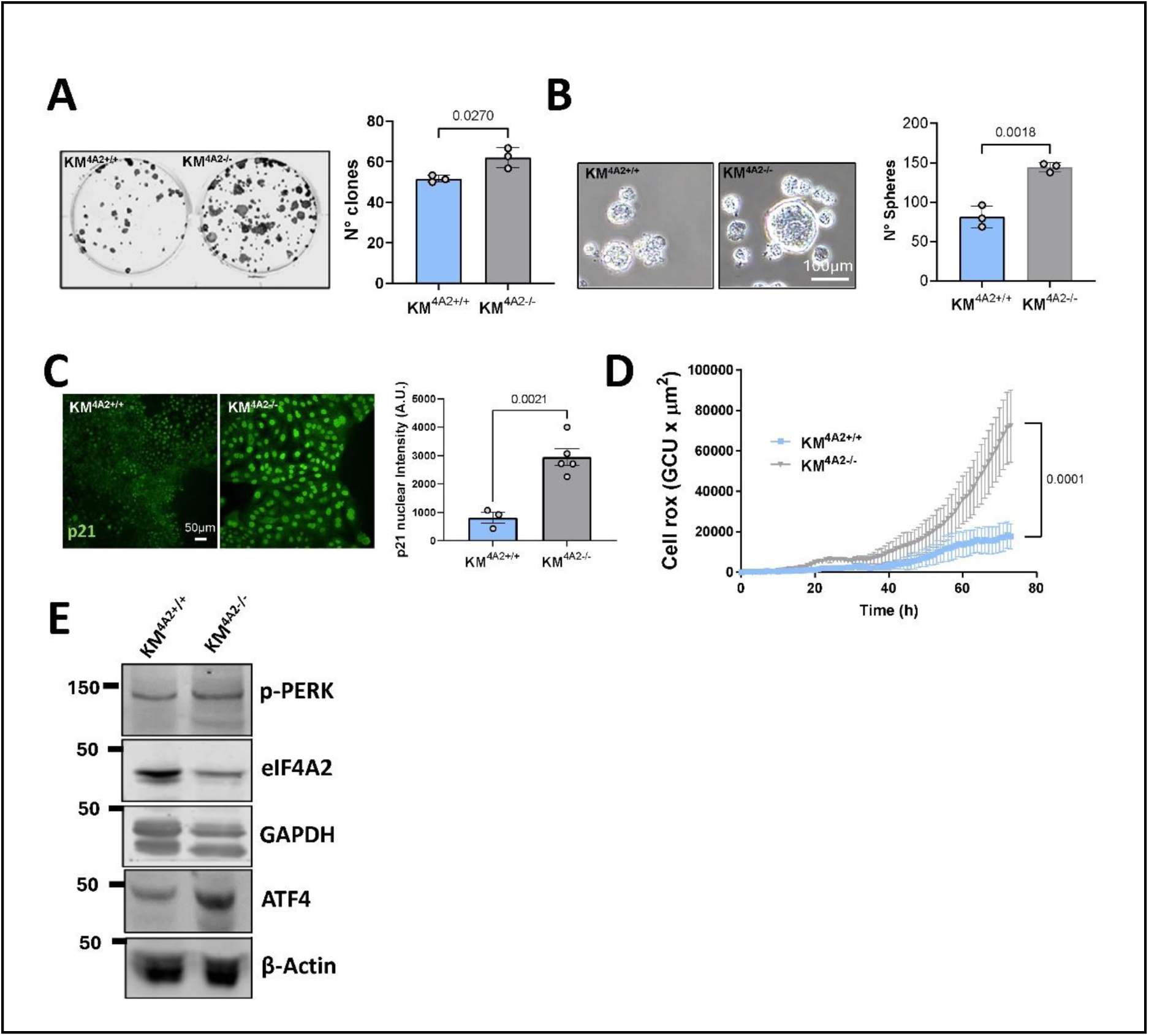
Phenotypic analysis of LKM^A2+/+^ and LKM^A2−/−^ cells. **(A,B)** KM^A2+/+^ and KM^A2−/−^ cells were plated at low density onto plastic dishes in serum containing medium **(A)** or into low adhesion wells in serum-free medium **(B)**. The resulting number of colonies **(A)** and tumour spheres **(B)** respectively were then determined. Values are mean ± SEM, N=3 individual experiments, statistical test is unpaired t-test. **(C, D)** KM^A2+/+^ or KM^A2−/−^ cells were plated onto plastic surfaces and allowed to grow for 48 hr. p21 expression was visualised and quantified using immunofluorescence **(C).** Cellular reactive oxygen species was determined using CellROX reagent **(D)**. Values are mean ± SEM, statistical test is unpaired t-test. **(E)** Activity of unfolded protein response signalling in KM^A2+/+^ and KM^A2−/−^ cells was assessed using Western blotting with antibodies recognising phospho-PERK and Atf4. β-actin and GAPDH were used as loading controls.

## Notes

### Competing Interest Statement

The authors have declared no competing interest.

